# Phage efficacy in infecting dual-strain biofilms of *Pseudomonas aeruginosa*

**DOI:** 10.1101/551754

**Authors:** Samuele Testa, Sarah Berger, Philippe Piccardi, Frank Oechslin, Grégory Resch, Sara Mitri

## Abstract

Bacterial viruses, or phage, play a key role in shaping natural microbial communities. Yet much research on bacterial-phage interactions has been conducted in liquid cultures involving single bacterial strains. Critically, phage often have a very narrow host range meaning they can only ever target a subset of strains in a community. Here we explore how strain diversity affects the success of lytic phage in structured communities. In particular, we infect a susceptible *Pseudomonas aeruginosa* strain PAO1 with lytic phage Pseudomonas 352 in the presence versus absence of an insensitive *P. aeruginosa* strain PA14, in liquid culture versus colonies growing on agar. We find that competition between the two bacterial strains reduces the likelihood of the susceptible strain evolving resistance to the phage. This result holds in liquid culture and in colonies. However, while in liquid the phage eliminate the whole sensitive population, colonies contain refuges wherein bacteria can remain sensitive yet escape phage infection. These refuges form mainly due to reduced growth in colony centers. We find little evidence that the presence of the insensitive strain provides any additional protection against phage. Our study reveals that living in a spatially structured population can protect bacteria against phage infection, while the presence of competing strains may instead reduce the likelihood of evolving resistance to phage, if encountered.

## Introduction

Lytic bacteriophage, or simply “phage”, are viruses that infect bacterial cells, replicate within them and then lyse them to spread and infect new hosts. Lytic phage are major bacterial predators that are highly abundant in number and distribution, thereby playing a key role in regulating bacterial population dynamics (1). Despite this potential importance, phage are rarely considered in studies of natural bacterial communities, such as the human microbiome project, or the Earth microbiome project – although this is beginning to change (2–5).

Their ability to reduce bacterial populations has also been harnessed as a therapeutic method, in “phage therapy”, whereby specific phage targeting a given bacterial pathogen is administered to patients to eliminate infections (6, 7). As we struggle to find solutions to tackle the emergence of antibiotic resistance (8), phage therapy has experienced renewed interest as a possible replacement or complementary treatment to antibiotics.

Although our appreciation of the importance of phage biology is on the rise, the experimental systems used to study phage still limit our understanding of their ecology and evolution in natural environments (9). Phages are typically studied in liquid cultures in the laboratory using a single phage and a single bacterial strain at a time. On the other extreme of the spectrum, clinical studies have been performed where phage cocktails are administered to animal or human hosts (10–13). Given all the complexity that such environments bring, it is difficult to explain differences between the results of laboratory and clinical studies (10, 11, 14, 15). Knowledge at an intermediate scale of complexity is clearly missing. Here, we expand on typical laboratory methods to study two dimensions of environmental complexity that likely matter in real microbial ecology: the presence of other bacterial strains, and life in a spatially-structured environment.

Bacteria rarely live in clonal groups, but typically share their environment with different microbial strains and species in dense, surface-attached cell groups called biofilms. Natural communities such as the human microbiome, or soil communities are hugely diverse (16, 17), including a large repertoire of phages (3, 18–20). Each of these phages tends to be quite host-specific, killing only a narrow range of bacterial strains (but see (21)). When phage attack a given target strain, we can expect little collateral damage to surrounding strains, and may therefore be tempted to also expect infection of the target to be independent of community structure. However, the presence of insensitive strains has been found to alter treatment outcomes by affecting target strain survival. Indeed, Harcombe & Bull (22) have shown that competition with a co-inhabiting species could reduce the ability of the targeted sensitive strain to survive phage attack. Their study considered liquid cultures, however. Since then, it has been shown that the spatial organization of different bacterial strains and species within biofilms can drive social interactions and the evolutionary trajectories of bacterial communities (23, 24). Biofilm-associated bacteria also have a higher survival rate compared to planktonic bacteria (25), particularly when exposed to antibiotics and importantly, also to phage (26). More generally, phage population dynamics differ radically between liquid bacterial cultures and bacteria growing on solid surfaces (27).

Here we show that both of these factors – the presence of other strains, and spatial structure – separately and combined affect the outcome of phage predation on the pathogen *Pseudomonas aeruginosa*, and its ensuing population dynamics. In particular, we target *P. aeruginosa* strain PAO1 with Pseudomonas phage 352 to which it is sensitive, in the presence and absence of a second strain, *P. aeruginosa* PA14 that is insensitive to the phage. Since phage are so specific, we believe the choice of a closely-related phage-insensitive strain to be a realistic one. We compare the outcome for PAO1 in a well-mixed liquid environment and a structured biofilm (colony) growing on a solid agar surface.

We find that in liquid, competition between the two strains can reduce the population size of the target strain PAO1, giving a competitive advantage to the phage and eliminating PAO1 without the emergence of resistance. Indeed, evolving resistance to the phage was the only way for PAO1 to survive phage attack in liquid. In contrast, in a biofilm treated with phage, PAO1 survived in the presence of the phage-insensitive strain PA14 without becoming resistant itself. Survival in the face of a phage attack, however, did not depend on PA14 but occurred in all biofilms, regardless of the presence of other strains. Instead, slower growth in the colony center appears to be the main mechanism that reduces the ability of the phage to replicate and spread through biofilms containing sensitive bacteria. The main effect of PA14 in the biofilm was instead in greatly reducing the likelihood of phage-resistance in PAO1.

## Results

### Inter-strain competition increases phage infectivity and reduces resistance evolution in liquid

We first sought to understand how treating a target strain *P. aeruginosa* PAO1 (henceforth PAO1) with Pseudomonas phage 352 in well-mixed liquid cultures is affected by the presence of a phage-insensitive strain *P. aeruginosa* PA14 (henceforth PA14). These liquid experiments involved growing bacteria in 96-well plates containing TSB and inoculated with mixtures of bacteria and phage over a period of 48 hours.

In control treatments with PAO1 growing alone, we observed that phage treatment resulted in a drop in PAO1 population size after 6 hours, after which the population recovered somewhat but not entirely (Fig. 1A). Assays testing for phage resistance (see Methods) revealed that after 24 hours of culture, 62 out of 63 tested colonies (98.41%) were resistant to the phage, while after 48 hours, 24 out of 24 (100%) were resistant. As a control, resistant PA14 cells growing alone were not significantly affected by the phage (Fig. 1B).

**Fig. 1.**
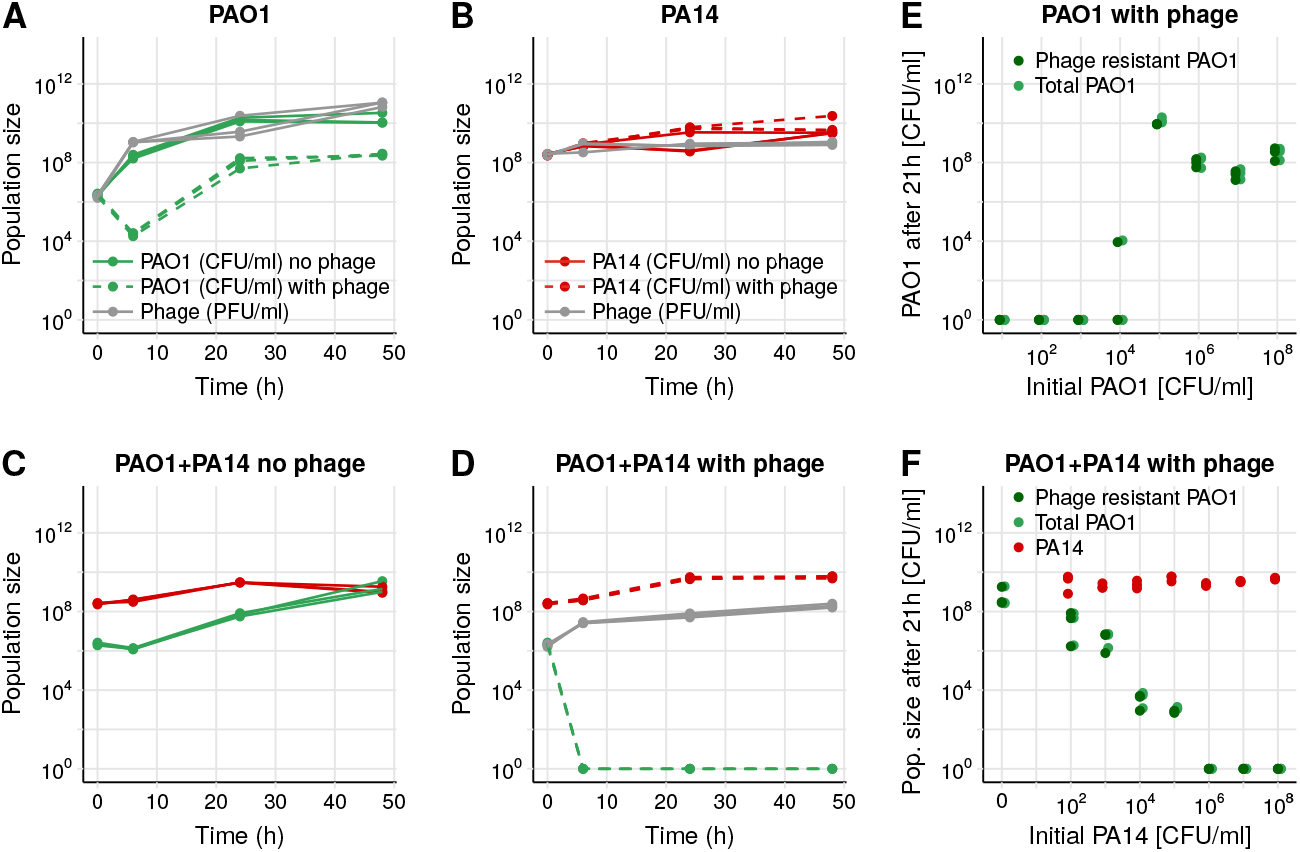
Phage efficacy in liquid. (A) Growth of PAO1 (in CFU/ml) in liquid over 48h. PAO1 grows without phage (solid green lines), but in its presence (dashed green lines) PAO1 decreases then rebounds, resulting in a resistant population (statistics in main text). The phage population (in PFU/ml, gray lines) increases accordingly. (B) PA14 (phage resistant), grows similarly in the presence or absence of phage (dashed or solid red lines, respectively), while phage (gray) remain approximately constant. (C) When PAO1 (green) and PA14 (red) are grown in co-culture in the absence of phage, PAO1 grows worse than alone. (D) When phage are added to the coculture, PAO1 population size drops below the detection limit at 6 hours and does not recover. (E) PAO1 is grown together with phage in triplicate at different initial population sizes (MOI=1). At the end of the experiment, bacteria are plated onto agar plates saturated with phage or not to count the resistant and total population (see Methods). A starting population size greater than ~10^4^ allows resistance to emerge. (F) Initial population size of PAO1 was always ~10^6^, while initial PA14 numbers varied as on the x-axis. Once PA14 became too numerous (greater than ~10^6^), PAO1 could no longer maintain its population size high enough for resistance to the phage to emerge. Red, light green and dark green points show population size of PA14, total PAO1 and resistant PAO1, respectively, at 21h.

Next, we co-cultured the two strains in the absence of phage and found that PAO1 grew worse than when it was alone, presumably due to competition with PA14 (Fig. 1C). Finally, adding the phage to this co-culture eliminated all PAO1 within 6 hours (Fig. 1D). Compared to growing alone then, PAO1 resistance could not emerge when growing with a competitor.

We hypothesized that the presence of PA14 prevented PAO1 from increasing its population size, thereby decreasing its potential to evolve resistance to the phage and survive the treatment. To test for the effect of population size on resistance evolution, we conducted two experiments. First, we grew PAO1 in the presence of phage with different starting population sizes, while maintaining the multiplicity of infection (MOI) constant at 1 (1 phage for each bacterium). In agreement with our hypothesis, resistance to the phage emerged when the initial population size was greater than 10^4^ CFU/ml (Fig. 1E). Second, we kept the initial population size of PAO1 constant at 10^6^ CFU/ml and varied the starting population size of its competitor PA14 in the presence of phage (MOI=1). Again, as predicted, phage resistance could emerge when there were fewer competitors, but once the number of competitors at the start exceeded 10^6^ CFU/ml, PAO1 cells were all killed by the phage at the end of 21 hours of co-culture (Fig. 1F). In all cases, PAO1 survival depended on becoming resistant to the phage.

In sum, in liquid culture, competition with a resident strain can prevent a targeted strain from surviving phage treatment, which is consistent with previous research (22).

### Phage infect sensitive PAO1 in mono-culture colonies

To simulate a setup where a biofilm forms on a solid surface and is later exposed to phage, we first grew the bacteria on a membrane filter placed on LB agar for approximately 12 hours until they had formed a small colony. We then transferred the filter with the 12-hour colony onto a new LB agar plate containing an air-dried drop (approximately the diameter of the filter) of either ~10^6^ phage, ~10^9^ phage, or no drop as a control. All colonies were left to grow in the presence or absence of the phage for an additional 36 hours, approximately (see Methods, Fig. 2A).

**Fig. 2.**
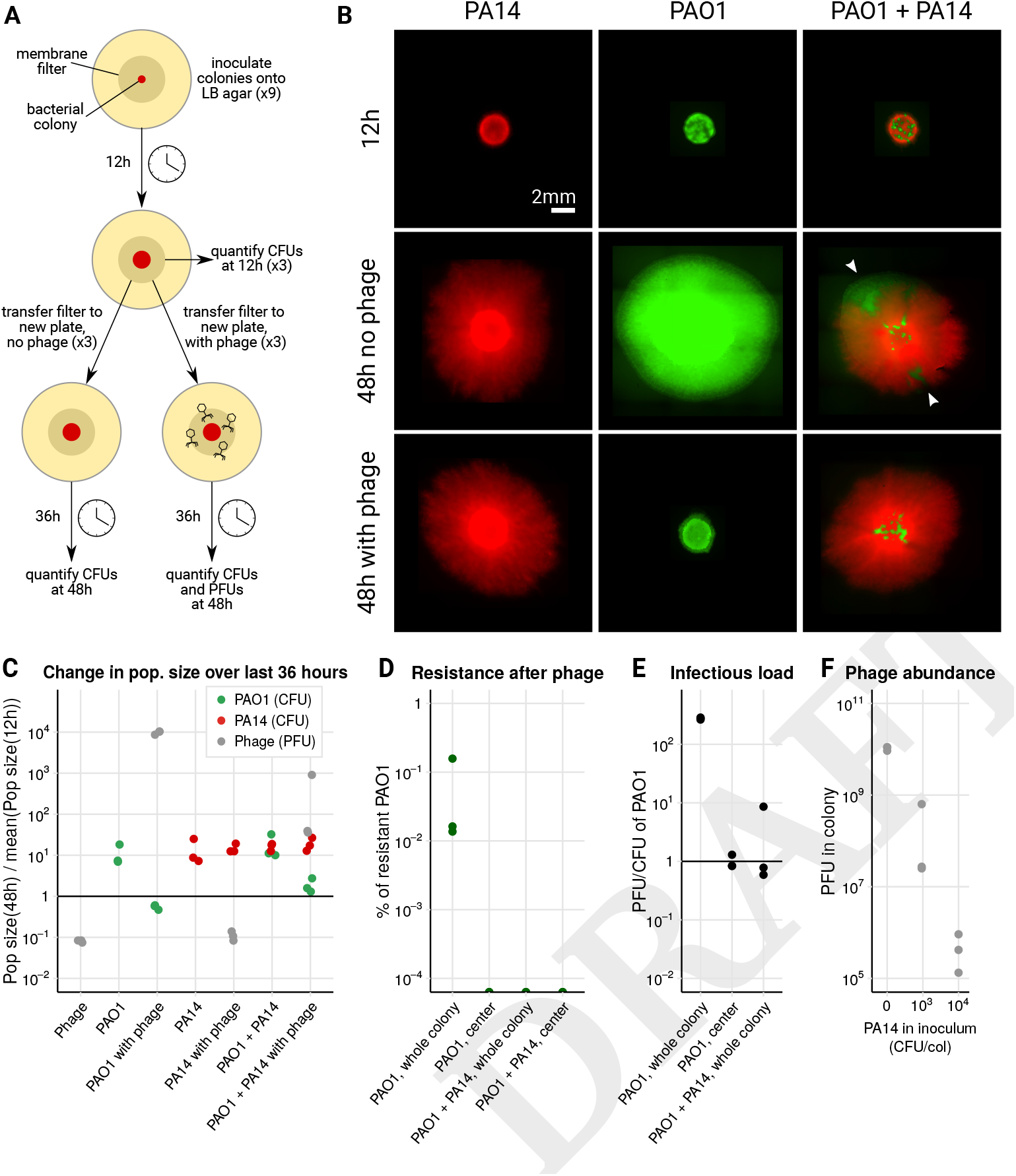
Phage efficacy in colonies. (A) All colonies(PAO1 alone, PA14 alone orPAO1 + PA14) were first grown by inoculating a drop of cells onto a membrane filter placed on a 0.1x LB agar plate in 9 replicates. After 12 hours (duration varied somewhat between experiments), 3 filters were removed to quantify CFUs, 3 were transferred to a new 0.1x LB agar plate containing a dried *50μl* drop of phage containing ~10^6^ PFU/ml, and 3 to a new 0.1x LB agar plate containing no phage. After 36 hours (with some variation) the remaining 6 filters were harvested to quantify CFUs and/or PFUs (see Methods). (B) Fluorescence microscopy images of the colonies at 12 and 48 hours. PA14 are tagged with mCherry (red) and PAO1 with GFP (green). Sectors that normally formed in untreated colonies (white arrowheads, center right) were absent in the phage treatment (bottom right) suggesting that phage kill cells at the colony edges where cells are most active, while cells in the center survive. See Fig. S3 for images from a similar experiment. (C-F) Data coming from triplicate colonies of a single experiment using unlabelled PA14. These data do not correspond to the images in (B), whose quantification was less precise (see Fig. S6) because PA14-mCherry were difficult to distinguish from PAO1-GFP (identical drug resistances). (C) the ratio of population sizes at 48 and 12 hours (see Fig. S1 for growth curves). Phage decrease the population size of PAO1 (green), resistant PA14 grow similarly across conditions (red), and phage decrease in the absence of PAO1 and increase in its presence. (D) We observed no phage resistance in the sampled centers of PAO1 colonies, or in colonies mixed with PA14. (E) To determine whether phage could reach colony centers, we quantified PAO1 and phage in the colony centers. The ratio of PFUs to CFUs was lower in the colony centers, and in whole mixed colonies compared to whole colonies of PAO1 alone. (F) Phage abundance in the colony at 48h is inversely proportional to the initial abundance of PA14 in the colony inoculum. PAO1 was held constant at 10^2^ CFU/col.

In PAO1 mono-culture colonies treated with phage, populations ceased to grow following phage arrival (comparison of CFUs at 12 and 48 hours, *df* = 29, *P =* 0.58, Fig. 2C, S1, S2), and there were significantly fewer bacteria in the phage treatment compared to the control (7.96±6.02×10^7^ with phage, versus 7.51±3.82×10^8^ without, *df* = 23, *P* < 0.001, Fig. S1, S2). Fluorescence microscopy images taken immediately prior to infection and 36 hours later showed that colonies treated with phage were smaller in diameter than non-treated colonies, with the fluorescent cells still visible in the center of the colony (Fig. 2B, middle column, Fig. S3). In the colonies that had been treated with phage, resistance to the phage was detected in 14 out of 15 colonies across five similar experiments, with resistant cells forming between ~0.04 and 20% of the total population at low (~10^6^ PFU/ml) initial phage dose (Fig. 2D, Fig. S5). At high initial phage dose (~10^9^ PFU/ml), the majority of surviving cells were found to be resistant to the phage, but a sub-population of sensitive cells survived in all replicates (Fig. S2F).

### Infection and the emergence of phage-resistance in PAO1 occurs mainly at colony edges

We wondered why so many sensitive cells survived and where in the colony resistance had occurred. To answer this question, before harvesting the colonies for quantification, we touched an inoculation loop in the center of the colony, resuspended its contents in PBS and plated the suspension to quantify the number of resistant and sensitive cells, as well as phage (see Methods). We found no resistant cells in the center of any of the colonies (Fig. 2D, Fig. S5), suggesting that resistance arose closer to the colony edges where most cellular growth occurs (28). Phage were nevertheless detected in the center, but the ratio of phage to uninfected cells was significantly lower than in the colony as a whole (PFU/CFU of 0.99±0.27 in the center and 276.5±11 in the whole colony, paired t-test, P<0.001, Fig. 2E). This suggests that the phage could spread to the center of the colony, but left a proportion of cells uninfected. Further evidence that some cells in the center were infected was that after washing to remove phage and plating on fresh agar, most cells lysed (Fig. S7), resembling “pseudolysogeny” or “hibernation”, which occurs in starved cells in stationary phase or persister cells, where phage DNA accumulates in the cell. Only once bacteria start growing again do viral capsids form and the phage resume their lytic cycle (27, 29–33). However, transmission electron microscopy images of the colony revealed many intact cells containing phage with assembled capsids that had not yet lysed, in addition to to some debris from lysed cells (Fig. S8). While phage were able to assemble – contrary to expectations for pseudolysogeny – the presence of unlysed and uninfected cells suggests a delay in lysis, which may explain why phage could not spread further and increase their numbers in the colony center.

### Phage penetration into colonies of insensitive PA14 is limited

In contrast, PA14 (the phage insensitive strain) mono-culture colonies were indistinguishable with and without phage treatment (Fig. 2B, C, t-test CFUs with and without phage, df=2.6, P=0.87, Fig. S1, S6). On sampling the colony centers, we never found phage in any of the colonies treated with a low phage dosage, but detected a few at the high initial dose of phage (on average 1 phage to every 863 PA14 cells). This suggests that phage could not diffuse much from the agar into PA14 colonies. Indeed, total phage populations fell to 11±2.8% of their original size in PA14 colonies over the 36 hours, which we suspect is due to toxicity of LB to phage (34) or temperature sensitivity, given that phage populations also fell to 8.1±5.4% in the absence of any bacteria (Fig. 2C, t-test with and without PA14: P=0.21). To determine whether phage could attach to PA14 cells, we performed an adsorption assay in liquid, and found that after 5 minutes of exposure to bacteria, phage only attached to PAO1 cells, but not PA14 (Fig. S9).

Taken together, in single-strain colonies we observe that PAO1 death and the emergence of phage resistance occurs mainly at the edges of the colony where cells are more actively growing. Only very few phage could spread into insensitive PA14 colonies at high phage titer, demonstrating that physical diffusion into colonies is very limited. Instead, cycles of attachment, infection and lysis allow phage to propagate deeper into colonies of sensitive PAO1. Phage can therefore infect some, but not lyse all PAO1 cells at the colony centers, where they are less metabolically active.

### Phage infect sensitive PAO1 in mixed colonies

Knowing that phage cannot diffuse much into PA14 colonies, we next asked how the presence of this insensitive strain would impact the survival of the targeted PAO1 within a colony containing both strains and treated with phage. We repeated the experiment (Fig. 2A) with a mixture of both PAO1 and PA14 at an initial ratio of 1:10, such that an approximate 1:1 ratio was reached on phage exposure after 12h (Fig. S1).

As in the phage-treated PAO1 mono-culture colony, PAO1 in the treated mixed colonies did not increase significantly following phage treatment (*df* = 17, *P* = 0.47, Fig. 2C, Fig. S2), and the phage treatment significantly reduced PAO1 cells compared to the untreated control (*df* =17, *P* < 0.001), demonstrating significant bacterial infection by phage. In addition, microscopy showed that patches of PAO1 (white arrowheads in Fig. 2B, right center) were absent from the edges of the colonies treated with phage (Fig. 2B, bottom right). Together, these data suggest that as in PAO1 colonies, cell lysis occurs at the actively growing edges.

We observed two significant differences between mono- and co-culture colonies, however. First, as in the liquid coculture, phage resistance was much less likely to emerge, with no phage resistance in mixed colonies infected with the low phage dose (Fig. 2D, S5), while at high infective dose, 0.6±0.3% of cells were resistant compared to the vast majority in the mono-culture colonies (Fig. S2). Second, co-culture colonies contained a lower infectious load (fewer phage per sensitive bacteria) compared to monoculture colonies at the end of the experiment (*df* = 20, *P* < 0.05, Fig. 2E, Fig. S2), indicating that phage could replicate less in the presence of PA14. To further verify this, we increased the number of PA14 in the colony inocula while keeping PAO1 constant, and found that phage abundance in the colony follows a strong negative correlation with initial PA14 abundance (Spearman’s *ρ* = 0.91, *P* < 10^−7^, Fig. 2F). Both these findings can be explained by what we observe in the images: since only a small proportion of the edge of a mixed colony is made up of PAO1 cells (Fig. 2B, white arrows), the effective population size of PAO1 (i.e. number of growing cells) is smaller in the presence of PA14 than in its absence (28, 35), making the emergence of resistance less likely (Fig. 1D), and keeping phage populations that infect them smaller (Fig. 2F).

Given that the phage seems to mainly infect PAO1 in the edges of both mixed and mono-culture colonies, we hypothesized that we would find phage refuges containing uninfected cells in both conditions, regardless of the presence of PA14.

### All colony centers contain phage-free refuges where sensitive PAO1 remain uninfected

To search for uninfected PAO1 in different areas of the colonies, we sampled the mono-culture and mixed colonies previously exposed to ~10^6^ PFU/ml by touching them with sterile toothpicks in four different locations (Fig. 3A), resuspending the cells and phage on the toothpicks in PBS and then spotting a drop of each suspension onto different agar plates to quantify the density of PAO1, phage, and PAO1 cells that had become resistant to phage (see Methods). To analyze these data (Fig. S10, S11), we imaged each drop after 15h of growth and processed the images (Fig. 3B) to quantify the density of healthy PAO1 cells and phage plaques in each position.

**Fig. 3.**
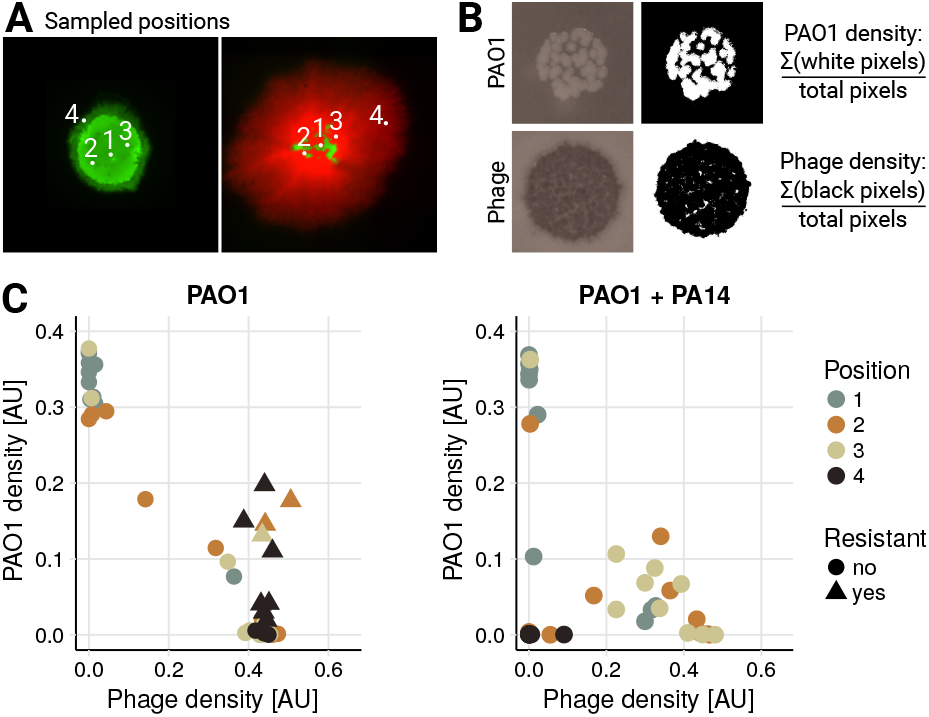
Sampling colonies to determine co-occurrence patterns of phage and bacteria. (A) The white dots indicate where we sampled in each colony using sterile toothpicks (same images as in Fig. 2B). Position 2 and 3 are equidistant from the center (position 1). (B) The toothpick-attached cells and phage were resuspended and a small drop of the suspension placed on LB agar containing gentamicin, and on soft LB agar containing PAO1. On the left are two representative images of these drops after ~15h at 37° C. To quantify the density of bacteria and phage, we applied a threshold to the images (two images on the right) and then calculated the proportion of white and black pixels in each picture respectively. These values are plotted in panel C. (C) Density of PAO1 and phage in each sample. Each dot or triangle corresponds to a sample in one position in one colony. The left and right panels show samples taken from 10 PAO1 and 10 mixed colonies, respectively (4×10=40 points on each plot). Resistance was determined by similarly thresholding images of drops grown on LB agar with gentamicin and saturated with ~10^10^ phage (see Fig. S10, S11 for the full data set). The different colors represent the positions sampled as shown in panel A.

In agreement with previous experiments, we only observed resistant PAO1 cells in samples coming from the monoculture colonies, and these were detected in positions sampled further away from the colony center (Fig. 3C, S12, triangles). Moreover, in line with our previous observation that PAO1 at the edge were killed by phage, sampling at the edge of the mixed colonies yielded very few PAO1 cells, and also very few phage (Fig. 3C, S12 black dots close to origin in right panel).

For all remaining samples (where sensitive PAO1 and phage density were > 0.1), phage density correlated negatively with the density of PAO1 (Pearson’s *ρ* = −0.9, *P* < 10^−10^), as one would expect. We also found that 35% of the samples from the mono-culture colonies and 20% from the mixed colonies had sensitive PAO1 cells close to the center that were completely uninfected by the phage (top left points in Fig. 3C, with PAO1 density > 0.25). This supports the presence of phage-protected refuges in the centers of all colonies, and rejects the hypothesis of phage-free refuges being caused solely by the presence of PA14.

This assay can be seen to represent a scenario where cells would have a chance to leave a biofilm and reseed a new environment. Cells from the refuges that were uninfected by phage would begin to grow and start new, healthy colonies (see also Fig. S7).

### Forced growth arrest and competition with a resistant PAO1 strain recapitulate our findings

To explain why many sensitive cells in the colony centers remained uninfected with or without PA14, we put forward two hypotheses that we tested next: first, that cells in the center of any colony can avoid phage infection because of a lack of growth; and second, that phage-resistant cells (not only insensitive PA14, but also newly emerging resistant cells) could create phage-free refuges in colonies by preventing phage spreading through reduced phage amplification. Accordingly, we repeated the experiment of Fig. 2A with two new conditions: in the first, we used phage-sensitive PAO1 cells (wildtype) but after the 12 hours of growth, we moved them onto agar lacking LB and containing EDTA to arrest bacterial growth, forcing them into stationary phase (36); and in the second we combined our wild-type PAO1 with 10 × more of a phage-resistant PAO1 strain isolated from the experiments described above (see Methods, Fig. S7F). These resistant mutants were found to be lacking the galU gene (Fig. S13), resulting in a loss of LPS and preventing phage attachment, as observed in previous work (37, 38) and verified by an adsorption assay (Fig. S9). A significant fitness cost was associated with the loss of LPS, as shown in a co-culture of wild-type and mutant PAO1 without phage (Fig. S15B, G).

The growth-arrested colony grew slightly (by 131±33.4% over 36 hours), and the phage increased 19.9-fold, approximately 3 orders of magnitude less than in a PAO1 colony growing on LB agar (Fig. S14A, E). Even though the phage replicated, they were not detected in the colony centers (0 in all three replicates). In contrast, phage were found in the centers of colonies of the mixed phage-sensitive and -resistant PAO1, at an infectious load that was similar to the monoculture colonies (Fig. S15J). In other words, even though the colony started with 10× more resistant cells, phage could still easily infect the sensitive bacteria and spread through the colonies (Fig. S15A, C). It is therefore unlikely that a rare mutant arising in a wild-type colony would provide protection to the sensitive cells, at least in part due to their reduced fitness. No phage were detected in the centers of control colonies containing only resistant PAO1, which were instead comparable to PA14 mono-culture colonies (Fig. S14F).

The data from these treatments support our previous conclusion: to reach the center of *P. aeruginosa* colonies, Pseudomonas phage 352 needs to attach to the surfaces of bacterial cells, and infect them while they are actively growing and dividing. Since phage-free refuges were observed in some mono-culture colonies where no resistance was detected (Fig. S11), and since phage could easily infect wild-type PAO1 in the presence of a large initial population of resistant PAO1, we conclude that growth arrest plays a more important role in protecting sensitive bacteria against phage infection compared to being surrounded by resistant or insensitive cells that phage cannot attach to or infect.

## Discussion

In sum (Fig. 4), we show that phage-sensitive bacteria are more likely to survive a phage attack if they are growing on a solid surface than in liquid culture. This is mainly because cells grow more slowly in the colony center, making the phage replicate more slowly and reducing their ability to amplify, lyse cells and spread into the center (Fig. 4B). The uninfected, phage-sensitive cells that remain can then potentially seed new, healthy bacterial colonies, if dispersed. We found little evidence that an insensitive strain (or a newly emerging resistant strain) further protects sensitive bacteria from the phage (Fig. 4C). Sensitive bacteria are, however, most likely to develop resistance to the phage in the absence of competing strains, where they can grow to a sufficiently large effective population size (Fig. 4B). Competitors thus reduce the likelihood of resistance evolution.

**Fig. 4.**
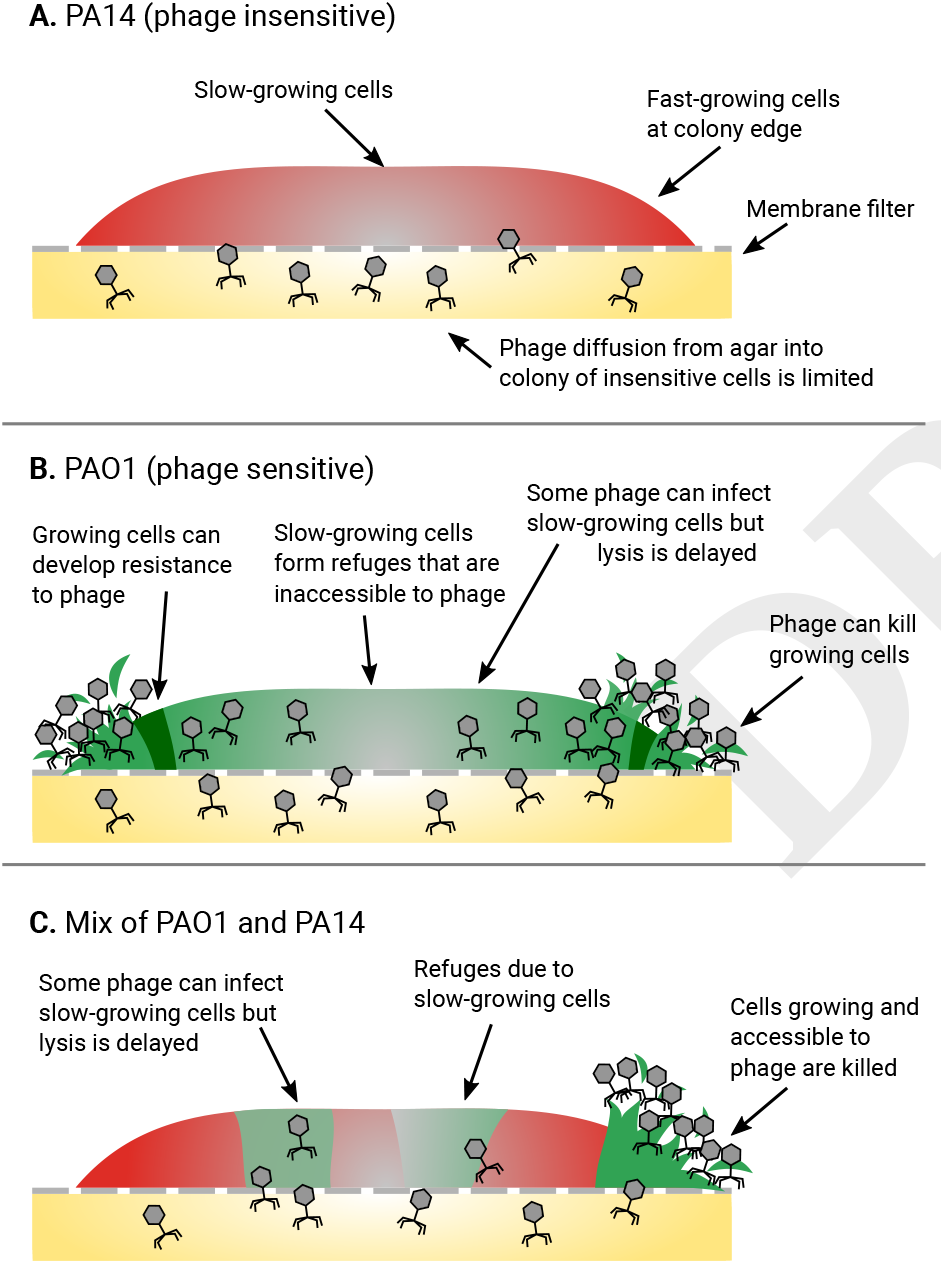
We propose a model for how Pseudomonas phage 352 infects colonies of single and mixed PAO1 and PA14 strains. Each drawing shows a cross-section of a bacterial colony, where higher bacterial growth rates are represented by solid colors and slower-growing bacteria by grey. (A) Phage infection and penetration into colonies of insensitive PA14 is limited. The same was found for PAO1 that had acquired resistance. (B) PAO1 colonies are increasingly infected towards the colony edges, correlating with growth rate (28). Phage resistance (dark green) is observed closer to the edges where growth and infection are occurring. Slow-growing cells toward the colony center form phage refuges. There, phage infect some cells of which only a subset is lysed. (C) In a mixture of sensitive and insensitive bacteria, insensitive cells reduce phage abundance overall, but phage-free refuges are mainly due to slow growth in the center. The emergence of phage resistance is limited in the presence of PA14.

Others have previously observed that sensitive bacteria can survive phage attack in biofilms (26, 39), and proposed that this may be due to a high bacterial density or large molecules that reduce phage diffusion, such as exopolysac-charides (40, 41). A recent study showed that *Escherichia coli* produces an amyloid fiber network that protects cells in a biofilm individually and reduces phage diffusion (41). Survival in the face of phage can instead occur because the bacteria reduce the expression of their phage receptors (42), or because they slow down growth as nutrients are depleted (40, 43, 44). Our data support a model whereby growth arrest can prevent phage infection (26, 32, 45–48).

We were curious whether spatially organized bacterial strains could protect each other against phage, as for different forms of environmental assault. For example, antibiotic-resistant cells can protect sensitive ones from antibiotics. This is because resistant cells can break down antibiotics and “detoxify” their local environment, allowing targeted susceptible cells in close proximity to survive and grow (49–52). Bacteria can also protect others against predators (53). To our knowledge, two studies have investigated cross-protection by infecting co-cultures of resistant and susceptible bacteria with phage. In the first, Tzipilevich *et al*. (54) found that rather than cross-protection, a sensitive strain of *Bacillus subtilis* actually conferred temporary phage-sensitivity to its previously resistant neighbor. This happened through the horizontal transfer of phage attachment molecules from lysed sensitive cells to intact resistant ones. In contrast, Payne *et al*. (55) demonstrated that cross-protection against phage T7 can occur between two strains of *E. coli*, where one harbored a CRISPR-based resistance. Cross-protection was observed both in liquid and on a bacterial lawn. One key difference to our study is that their CRISPR-immune cells remove the phage from the environment through adsorption and degradation, and then stop growing, whereas in our system, phage do not even attach to insensitive PA14 cells. PA14 cells simply do not seem to interact with the phage.

The range of different experimental outcomes leads us to conclude that cross-protection against phage strongly depends not only on growth conditions, but also on the choice of phage and bacteria, including their resistance mechanism. First, our finding that Pseudomonas phage 352 cannot infect growth-arrested sensitive cells may not apply to other phages. Phage T4, for example, is similarly unable to infect stationary phase *E. coli* cells, but phage T7 can (33, 56). Other phages of different sizes or hydrophobicities may be better or worse at diffusing through biofilm (57). Second, it is possible that a greater protective effect between our particular strains would be observed under different growth conditions, for example if we were to provide our bacteria with nutrients that were constantly replenished to limit growth arrest. Third, since PAO1 and PA14 compete with one another, they tend to separate in space. Strains that rely on each other to grow have instead been shown to remain mixed in colonies (58, 59). Increased mixing may then increase cross-protection against phage.

Collectively, we are revealing that the way in which phage shape microbial community dynamics depends on strain identity and environmental conditions. This realization may explain why in phage therapy, for example, such large discrepancies are observed between laboratory results and *in vivo* trials (10, 11, 14, 15). But despite the simplicity of our colony model relative to an *in vivo* system, our insights are important: Biofilms - a typical mode of growth in an infection - appear to be more difficult to treat with phage compared to liquid cultures; a fitness trade-off between phage resistance and fitness (Fig. S15) is likely to be very common, meaning that phage-resistant strains are typically less virulent than wildtype strains (reviewed in (13)). Resistance evolution is therefore likely to pose less of a threat in practice than cells remaining completely uninfected by phage and dispersing to cause new phage-free infections. We still need to understand whether and under what conditions strain diversity can reduce phage efficacy, but here we show that competition with other strains can reduce the likelihood of resistance evolution. Future research should go beyond dual-strain interactions to better mimic the diversity in natural environments.

Finally, we highlight the importance of spatial structure for the ecology and evolution of microbial populations. In a liquid environment, phage may drive sensitive strains locally extinct, potentially destabilizing the bacterial community. In a multi-strain biofilm, phage may instead generate diversity through uneven infection, which creates local areas of either phage-resistant, phage-infected or phage-protected bacteria (Fig. 3C, Fig. 4), each subject to different selection pressures (43). In turn, phage have access to different bacterial niches (43, 60). The resulting co-evolutionary dynamics mean that spatially organized bacteria-phage populations, which are likely to be the norm in many environments, may be key to maintaining the diversity, stability and the evolvability of microbial communities.

## Materials and Methods

### Bacterial strains, phage, media and culture conditions

Experiments were performed with two different strains of *Pseudomonas aeruginosa*: strain PAO1 modified with a miniTn7 transposon containing a GFP or DsRed marker, which was susceptible to a specific phage, and strains PA14 (PA14-WT) or modified with a Tn7 transposon containing an mCherry marker (PA14-mCherry), which were both resistant to this same phage. Both transposons contained a gentamicin resistance gene. All three strains were kindly provided by Kevin Foster. The phage used for this study was Pseudomonas phage 352, Myoviridae morphotype A1, previously *φ*14 (61, 62) (received from D. Haas, J.-F. Vieu, E. Ashenov and R. Lindberg). We chose this phage among 14 that we tested based on a spot assay that produced phage plaques on PAO1-GFP but not the two PA14 strains, which were entirely resistant.

Overnight cultures were grown in tryptic soy broth (TSB; Bacto™, Detroit, MI, USA) at 37°C, shaken at 200 rpm. Before each experiment, the optical density (OD_600_) of the overnight cultures of PAO1-GFP, and either PA14-mCherry or PA14-WT strains (depending on the experiment) was measured with a spectrophotometer (Ultrospec 10, Amersham Biosciences). Bacterial overnight cultures were then inoculated into Erlenmeyer flasks (100 ml) containing 20 ml of TSB to obtain a standardized OD_600_ of 0.05. Bacterial cultures were grown in a shaking incubator at 200 rpm and 37°C for 3 hours to obtain bacteria in exponential phase with a final concentration of approximately 10^8^ CFU/ml at the beginning of each experiment. These cultures were then diluted in PBS to the desired starting population size.

### Quantifying bacterial and phage populations

To quantify bacterial colony-forming units (CFU) and plaque-forming units (PFU) of phages, in liquid assays, CFUs and PFUs were measured directly, while for colonies, bacteria and phage (if applicable) were first extracted from the filters and suspended in PBS (see below). Suspensions coming from phage-treated liquid cultures or colonies were centrifuged at 8000 rpm for 15 minutes at 4°C. Following centrifugation, the supernatant containing phage was kept in the fridge at 4°C and later used to measure PFUs. The centrifuged bacterial pellet was resuspended in either 200μl (liquid) or 1ml (colonies) of PBS, and then washed 3 times with 1 ml of fresh PBS at 8000 rpm, 4°C for 5 minutes to remove all the potential phages remaining in the pellet.

CFUs were quantified by serially diluting all cell suspensions (from liquid cultures or colonies, with or without phage) from 10^0^ to 10^−7^ in PBS, and spreading 10μl drops in lines across Tryptic Soy Agar (TSA) or Luria-Bertani (LB) agar plates. After 15h in a 37°C incubator, colonies were counted at the most appropriate dilution. To distinguish the two *P. aeruginosa* strains, co-cultures were plated onto TSA or LB agar plates to count non-fluorescent PA14-WT CFUs and onto LB agar plates containing 10*μ*g/ml of gentamicin to count only PAO1-GFP CFUs. In experiments where PAO1-GFP and PA14-mCherry were co-cultured, both strains were resistant to gentamicin, and were only plated onto TSA or LB agar and distinguished by their fluorescence. PFUs were quantified similarly, except that drops were spread in lines across 20ml soft LB agar (30g/L TSB + 7g/L agar) mixed with 300μl of PAO1 overnight culture and allowed to dry for 1h in a laminar flow hood. For the treatments involving phage in colonies, the whole agar was also collected and put in 50 ml falcon tubes containing 10 ml of PBS, well-shaken, centrifuged for 15 minutes at 4000 rpm at 4°C and the supernatant containing phages further diluted in PBS to count the PFUs. Phage concentrations from the filter including the colony, the agar and the touched colony center (see below) were summed up to obtain the final PFU/colony value.

The rate of resistance of PAO1 to the phage was calculated in two ways. In the first method, 10*μl* of a solution containing 10^10^ PFU/ml was streaked in a straight line across an LB agar plate. Then, having previously plated PAO1 to count CFUs, individual colonies were picked and streaked in parallel lines perpendicular to the line containing phage, and the plate incubated for 15h at 37°C. Picked colonies that resulted in solid lines across the length of the plate were classified as resistant, while bacteria in lines that were truncated where the phage had been spread were considered to be sensitive. For the second method, we plated cultures on TSA plates on which we had previously spread 500*μ*l containing approximately 10^10^ PFU/ml of pre-absorbed phages, and allowed to dry. If PAO1-GFP were growing in co-culture with PA14-WT, plates additionally containing 10 *μ*g/ml gentamicin were used. To evaluate resistance rates, the CFUs/colony of PAO1-GFP growing on plates saturated with phages (resistant) was then compared to the CFUs/colony growing on plates with no phage (total uninfected).

### Phage treatment in liquid cultures

A 96-well plate was filled with 200 μl of TSB in each well, additionally containing 10^6^ CFU/ml PAO1-GFP or 10^8^ CFU/ml PA14-WT alone, or together with or without 10^6^ PFU/ml of phages (MOI(PAO1)=1). In PA14 mono-cultures, 10^8^ PFU/ml were inoculated (MOI(PA14)=1). Initial population sizes of bacteria and phage were quantified prior to mixing. Each condition (PAO1-GFP alone, PA14-WT alone, and the co-culture) was performed in triplicate. These initial population sizes were chosen since they allowed the two strains to co-exist and grow over 48 hours (PAO1 is more competitive). The plate was then put in a Tecan Infinite 200 PRO plate reader at 37°C under agitation for 48 hours. After 6, 24 and 48 hours, the samples were transferred into Eppendorf tubes, washed, serially diluted and plated as described above.

### Quantifying phage resistance rates in liquid

To understand the role of population size on resistance emergence, two experiments were performed (Fig. 1E, F). In the first, a 96-well plate was filled with 10 up to 10^8^ CFU/ml of PAO1-GFP, with 10-fold increases, together with phage to achieve an MOI(PAO1)=1 in 200 μl of TSB. We grew the bacteria for 21 hours at 37°C under agitation in the plate reader, and then assessed phage resistance rates and total population sizes as described above. For the second experiment, a 96-well plate was filled with 10^6^ CFU/ml of PAO1-GFP and phages at an MOI(PAO1)=1 in 200 μl of TSB, to which we added increasing amounts of PA14, starting at 10^2^ up to 10^8^ in 10-fold increments. Bacteria were again grown in the plate reader for 21 hours at 37°C under agitation, at the end of which we assessed phage resistance rates and total population sizes of both strains as described above.

### Colony experiments and phage treatment

To grow bacteria in a colony, liquid cultures were prepared and a drop spotted onto a membrane filter (Isopore^®^ Membrane, 0.2μm PC membrane, GTTP02500, Merck) previously placed in the centre of agar plates containing 0.1x LB (1 g/L of tryptone (ThermoScientific™ Oxoid™ Tryptone), 0.5 g/L of yeast extract (ThermoScientific™ Oxoid™ Yeast Extract Powder), 10 g/L of NaCl (ACROS Organics™, 99.5%) and 15 g/L of agar (Bacto™ agar solidifying agent, BD Diagnostics)). Liquid cultures of the two strains were prepared as described for the liquid experiments and diluted in PBS to obtain a final concentration of 10^4^ CFU/ml of PAO1-GFP and 10^5^ or 10^6^ CFU/ml of PA14-WT (for a ratio of 1:10 or 1:100, respectively). 100μl of each strain were mixed together, or with 100μl of PBS for the mono-culture colonies. A 2μl drop of the mixture was then spotted onto the filter. Nine replicate plates were prepared for each condition (PAO1-GFP, PA14-WT or mCherry, and the mixture of both), and incubated at 37°C. After 12 hours of incubation, three replicates were removed in order to count the CFUs of both strains by removing the filters from the agar using sterile tweezers and placing them in tubes containing 3 ml of PBS. The tubes were extensively vortexed to remove and resuspend the colonies in the PBS, the filters removed and the bacteria plated to count CFUs as described above. Among the six remaining replicates, three were placed onto new 0.1x LB agar plates without phage and the three others were placed onto new 0.1x LB agar plates pre-absorbed with a 50μl drop containing ~10^6^ or ~10^9^ phages (diameter similar to filter diameter) depending on the experiment, and incubated at 37°C. After ~36 hours, to quantify phage infectious load in the colony center of phage-treated colonies, we touched a sterile, plastic inoculation loop to the top center of each colony (without going deep enough to touch the filter) and resuspended its contents in 1ml of PBS. We then quantified CFUs and PFUs of this suspension as described above. The Isopore^®^ filters with the remaining majority of the colony were then carefully removed with sterile tweezers, resuspended in 3ml of PBS, and the suspension used to quantify CFUs and PFUs as described above (the phage-treated colonies with the centrifugation step described previously). Finally, the remaining agar was used to quantify PFUs as described above. For the experiments where we arrested the growth of PAO1 after 12 hours, agar plates were prepared containing 10g/L of NaCl, 15 g/L of Bacto™ agar and 0.05mM of EDTA. These plates were either spotted with a drop containing phage or no drop, and the filters transferred onto them as described above.

### Phage adsorption test

To test whether phage could adsorb to the different strains, we prepared (on ice) a solution containing ~10^6^ bacteria (either PAO1, PA14, PA14-mCherry or PAO1 res2) and added ~10^6^ phage to each. We quantified the PFUs in the starting inoculum of phage as described above. After 5 and 10 minutes on ice, we filtered 2ml of the suspension using 3ml Omnifix^®^ syringe filters with a pore size of 0.22μm (Cobetter^®^), and quantified the PFUs in the supernatant as described above. A reduction of phage in the supernatant indicated that the phage had attached to the cells, and ended up in the pellet rather than the supernatant (Fig. S9).

### Toothpick sampling assay

To assess where in the colonies phage and infected or uninfected PAO1 bacteria were located, the experiments were repeated using 10 replicate PAO1 colonies and 10 mixed (PAO1 and PA14) colonies. We defined four locations to sample from in each colony as shown in Fig. 3A, taking care to sample only from the top of that area (not going so deep as to touch the filter). Note that this is not a very precise method. Each toothpick was then suspended in 300ml of PBS, vortexed, and 5μl of the resulting solution inoculated onto (i) LB agar plates to quantify overall bacterial density, (ii) gentamicin-containing LB agar plates to quantify PAO1 density, (iii) gentamicin-containing LB agar plates saturated with approximately 10^10^ PFU/ml of pre-absorbed phages to quantify PAO1 resistance and (iv) TSB + soft agar containing PAO1 as described above to quantify phage density. After ~15 hours of growth at 37°C, we imaged each of the resulting spots using a Dino-Lite Edge microscope.

### Microscopy and image analysis

Images of the colonies were acquired after 12 and 48 hours using a Zeiss AXIO Imager M1 fluorescence microscope and a 2.5x objective. PAO1-GFP colonies were imaged using a GFP filter set (excitation: 470/40, emission: 525/50) and PA14-mCherry colonies using an mCherry filter set (excitation: 545/30, emission: 620/60, with automatic exposure), and for mixed colonies, an overlay of the two images was produced using imageJ. Since some colonies after 48 hours were too large to fit in one image, a series of 3× 3 images were acquired and stitched together using autostitch software (63). For the toothpick sampling assay, each image was manually cropped to 600×600 pixels, converted to grayscale, and a threshold applied using the Matlab Image Processing toolbox to yield the photos in Fig. 3B, S10, and S11. We then summed the black pixels and white pixels to compute phage or bacterial density, respectively, and divided them by the total number of pixels.

For the transmission electron microscopy (Fig. S8), the filter and colony were removed with sterile tweezers, placed upside-down and fixed in a 2.5% glutaraldehyde solution (EMS, Hatfield, PA) in phosphate buffer (PB 0.1 M, pH 7.4) for 1h at room temperature (RT) and post-fixed in a fresh mixture of 1% osmium tetroxide (EMS) with 1.5% of potassium ferrocyanide (Sigma, St. Louis, MO) in PB buffer for 1h at RT. The samples were then washed twice in distilled water and dehydrated in ethanol solution (Sigma, St Louis, MO, US) at graded concentrations (30% for 40 mins; 50% for 40 mins; 70% for 40 mins; 100% for 2×30 mins). This was followed by infiltration in 100% Epon resin (EMS, Hatfield, PA, US) over-night, and finally polymerized for 48h at 60°C in an oven. Ultrathin sections of 50nm thick were cut transversally to the filter, using a Leica Ultracut (Leica Mikrosystem GmbH, Vienna, Austria), picked up on a copper slot grid 2× 1mm (EMS, Hatfield, PA, US) coated with a polystyrene film (Sigma, St Louis, MO, US). Sections were post-stained with uranyl acetate (Sigma, St Louis, MO, US) 4% in water for 10 mins, rinsed several times with water followed by Reynolds lead citrate in water (Sigma, St Louis, MO, US) for 10 mins and rinsed several times with water. Micrographs were taken with a transmission electron microscope FEI CM100 (FEI, Eindhoven, the Netherlands) at an acceleration voltage of 80kV with a TVIPS TemCamF416 digital camera (TVIPS GmbH, Gauting, Germany).

### Identifying resistance mutation in PAO1

PAO1 mutant cells were added to 45 *μ*l of lysis buffer (10mM Tr-isHCL, 1mM EDTA, 0.1% Triton X adjusted to pH 8.0 using NaOH; 2.5*μ*l of 20mg/ml solution of lysozyme, Sigma-Aldrich, 62971-10G-F; 2.5μl of 10mg/ml proteinase K, Sigma-Aldrich). The sample was lysed using a thermocycler (20 mins at 37^°^C, 20 mins at 55^°^C, 20 mins at 95^°^C). The galU gene was amplified from the lysate with forward (5’-CCGACAAGGAAAAGTACCTGG-3’) and reverse (5’-CGCTTGCCCTTGAACTTGTAG-3’) primers. The reaction mixture (25 μl, final volume) contained 15.375 μl of nuclease-free water, 5 *μ*l of 5× Gotaq buffer (Promega M792A), 1μl of 10μM forward primer, 1ul of 10μM reverse primer, 0.5 μl of 10μM dNTP mix (Promega U151B), 1U of GoTaq G2 DNA polymerase (Promega, M784B) and 2*μ*l of bacterial lysate. The PCR was performed with the thermocycler: 5 mins of initial denaturation at 95°C, followed by 35 cycles of denaturation (30s at 95°C), annealing (30s at 55^°^C), and extension (50s at 72^°^C), with a final extension step (8 mins at 72° C). Amplified products from all samples were verified by gel electrophoresis (Fig. S13).

### Statistical analysis

Each experiment was performed using three biological replicates per condition. Due to this low replicate number, we compared treatments using two-tailed t-tests. Experiments were then repeated on separate occasions, and results are reported in supplementary material. We combined data from corresponding treatments across experiments by fitting a linear model to the data and applying a blocked ANOVA test. To test whether phage and bacterial densities correlated in the toothpick assay, we used Pearson’s correlation test.

## ACKNOWLEDGEMENTS

We would like to thank Harald Brüssow, Kevin Foster, Flor Arias-Sánchez and Shawna McCallin for insightful discussions. We thank Kevin Foster for the bacterial strains, Dieter Haas posthumously for the phage, Marc Garcia-Garcerà for extracting the DNA and running a PCR on the mutant strain, and Semhar Ghebre-hiwet Tekle for constructing the mCherry plasmid. We appreciate the assistance of Damien De Bellis and the Electron Microscopy Facility (EMF) at the University of Lausanne (UNIL) and thank them for their support. ST and PP were funded by the University of Lausanne, FO by an SNF Early Postdoc. Mobility grant, SB and SM by ERC Starting grant 715097.

## Bibliography

1. Lucía Fernández, Ana Rodríguez, and Pilar García. Phage or foe: an insight into the impact of viral predation on microbial communities. The ISME Journal, 2018. ISSN 1751-7362. doi: 10.1038/s41396-018-0049-5.

2. Mohammadali Khan Mirzaei and Corinne F. Maurice. Ménage à trois in the human gut: interactions between host, bacteria and phages. Nature Reviews Microbiology, 15(7):397–408, may 2017. doi: 10.1038/nrmicro.2017.30.

3. Pauline D Scanlan. Bacteria-Bacteriophage Coevolution in the Human Gut: Implications for Microbial Diversity and Functionality. Trends in microbiology, 393(0):16–23, mar 2017. doi: 10.1016/j.tim.2017.02.012.

4. Jennifer R et al Brum. Ocean plankton. Patterns and ecological drivers of ocean viral communities. Science, 348(6237):1261498, may 2015. doi: 10.1126/science.1261498.

5. Felipe H. Coutinho et al. Marine viruses discovered via metagenomics shed light on viral strategies throughout the oceans. Nature Communications, 8:15955, jul 2017. doi: 10.1038/ ncomms15955.

6. Stephen T. Abedon, Pilar Garcia, Peter Mullany, and Rustam Aminov. Editorial: Phage Therapy: Past, Present and Future. Frontiers in Microbiology, 8:981, jun 2017. doi: 10. 3389/fmicb.2017.00981.

7. Jeroen De Smet, Hanne Hendrix, Bob G. Blasdel, Katarzyna Danis-Wlodarczyk, and Rob Lavigne. Pseudomonas predators: understanding and exploiting phage-host interactions. Nature Reviews Microbiology, jun 2017. ISSN 1740-1526. doi: 10.1038/nrmicro.2017.61.

8. Evelina Tacconelli et al. Discovery, research, and development of new antibiotics: the WHO priority list of antibiotic-resistant bacteria and tuberculosis. The Lancet Infectious Diseases, 0(0), dec 2017. ISSN 14733099. doi: 10.1016/S1473-3099(17)30753-3.

9. Marta Lourenço, Luisa De Sordi, and Laurent Debarbieux. The Diversity of Bacterial Lifestyles Hampers Bacteriophage Tenacity. Viruses, 10(6), 2018. doi: 10.3390/v10060327.

10. C Lerondelle and B Poutrel. [Bacteriophage treatment trials on staphylococcal udder infection in lactating cows (author’s transl)]. Annales de recherches veterinaires. Annals of veterinary research, 11(4):421–6, 1980. ISSN 0003-4193.

11. Haiqing Sheng, Hannah J Knecht, Indira T Kudva, and Carolyn J Hovde. Application of bacteriophages to control intestinal Escherichia coli O157:H7 levels in ruminants. Applied and environmental microbiology, 72(8):5359–66, aug 2006. doi: 10.1128/AEM.00099-06.

12. R J Atterbury et al. Bacteriophage therapy to reduce salmonella colonization of broiler chickens. Applied and environmental microbiology, 73(14):4543–9, jul 2007. doi: 10.1128/ AEM.00049-07.

13. Frank Oechslin. Resistance Development to Bacteriophages Occurring during Bacteriophage Therapy Viruses, 10(7):351, jun 2018. doi: 10.3390/v10070351.

14. Damien Maura, Eric Morello, Laurence du Merle, Perrine Bomme, Chantal Le Bouguénec, and Laurent Debarbieux. Intestinal colonization by enteroaggregative Escherichia coli supports long-term bacteriophage replication in mice. Environmental Microbiology, 14(8):1844–1854, 2012. ISSN 14622912. doi: 10.1111/j.1462-2920.2011.02644.x.

15. Marine Henry, Rob Lavigne, and Laurent Debarbieux. Predicting in vivo efficacy of therapeutic bacteriophages used to treat pulmonary infections. Antimicrobial agents and chemotherapy, 57(12):5961–8, dec 2013. doi: 10.1128/AAC.01596-13.

16. Daniel Schlatter, Linda Kinkel, Linda Thomashow, David Weller, and Timothy Paulitz. Disease Suppressive Soils: New Insights from the Soil Microbiome. Phytopathology, 107(11): 1284–1297, nov 2017. doi: 10.1094/PHYTO-03-17-0111-RVW.

17. Peter J Turnbaugh, Ruth E Ley, Micah Hamady, Claire M Fraser-Liggett, Rob Knight, and Jeffrey I Gordon. The human microbiome project. Nature, 449(7164):804–10, oct 2007. ISSN 1476-4687. doi: 10.1038/nature06244.

18. Lesley A. Ogilvie and Brian V. Jones. The human gut virome: a multifaceted majority. Frontiers in Microbiology, 6:918, sep 2015. doi: 10.3389/fmicb.2015.00918.

19. Samuel Minot, Alexandra Bryson, Christel Chehoud, Gary D. Wu, James D. Lewis, and Frederic D. Bushman. Rapid evolution of the human gut virome. Proceedings of the National Academy of Sciences, 110(30):12450–12455, jul 2013. doi: 10.1073/PNAS.1300833110.

20. Michiel Vos, Philip J Birkett, Elizabeth Birch, Robert I Griffiths, and Angus Buckling. Local adaptation of bacteriophages to their bacterial hosts in soil. Science, 325(5942):833, aug 2009. doi: 10.1126/science.1174173.

21. Kathryn M. Kauffman and Martin F. Polz. Streamlining standard bacteriophage methods for higher throughput. MethodsX, 5:159–172, jan 2018. doi: 10.1016/J.MEX.2018.01.007.

22. W. R. Harcombe and J. J. Bull. Impact of phages on two-species bacterial communities. Applied and Environmental Microbiology, 71(9):5254–5259, 2005. ISSN 00992240. doi: 10.1128/AEM.71.9.5254-5259.2005.

23. Sara Mitri and Kevin Richard Foster. The Genotypic View of Social Interactions in Microbial Communities. Annual Review of Genetics, 47:247–73, aug 2013. ISSN 1545-2948. doi: 10.1146/annurev-genet-111212-133307.

24. Carey D. Nadell, Knut Drescher, and Kevin R. Foster. Spatial structure, cooperation and competition in biofilms. Nature Reviews Microbiology, 14(9):589–600, jul 2016. ISSN 1740-1526. doi: 10.1038/nrmicro.2016.84.

25. Philip S Stewart and J William Costerton. Antibiotic resistance of bacteria in biofilms. The Lancet, 358(9276):135–138, jul 2001. doi: 10.1016/S0140-6736(01)05321-1.

26. Rasmus Skytte Eriksen, Sine L Svenningsen, Kim Sneppen, and Namiko Mitarai. A growing microcolony can survive and support persistent propagation of virulent phages. Proceedings of the National Academy of Sciences of the United States of America, 115(2):337–342, jan 2018. doi: 10.1073/pnas.1708954115.

27. Stephen T. Abedon. Bacteriophage ecology: population growth, evolution, and impact of bacterial viruses. Cambridge University Press, 2008. ISBN 9780511541483.

28. Sara Mitri, Ellen Clarke, and Kevin R Foster. Resource limitation drives spatial organization in microbial groups. The ISME Journal, 10(6):1471–1482, jun 2016. ISSN 1751-7362. doi: 10.1038/ismej.2015.208.

29. T. A. Kokjohn and G. S. Sayler. Attachment and replication of Pseudomonas aeruginosa bacteriophages under conditions simulating aquatic environments. Journal of General Microbiology, 137(3):661–666, mar 1991. doi: 10.1099/00221287-137-3-661.

30. S. Ripp and R. V. Miller. The role of pseudolysogeny in bacteriophage-host interactions in a natural freshwater environment. Microbiology, 143(6):2065–2070, jun 1997. doi: 10.1099/ 00221287-143-6-2065.

31. Marcin Los, Grzegorz Wegrzyn, and Peter Neubauer. A role for bacteriophage T4 rI gene function in the control of phage development during pseudolysogeny and in slowly growing host cells. Research in Microbiology, 154(8):547–552, oct 2003. doi: 10.1016/ S0923-2508(03)00151-7.

32. Sivan Pearl, Chana Gabay, Roy Kishony, Amos Oppenheim, and Nathalie Q Balaban. Non-genetic Individuality in the Host-Phage Interaction. PLoS Biology, 6(5):e120, may 2008. doi: 10.1371/journal.pbio.0060120.

33. Daniel Bryan, Ayman El-Shibiny, Zack Hobbs, Jillian Porter, and Elizabeth M. Kutter. Bacteriophage T4 Infection of Stationary Phase E. coli: Life after Log from a Phage Perspective. Frontiers in Microbiology, 7:1391, sep 2016. doi: 10.3389/fmicb.2016.01391.

34. H. Hadas, M. Einav, I. Fishov, and A. Zaritsky. Bacteriophage T4 Development Depends on the Physiology of its Host Escherichia Coli. Microbiology, 143(1):179–185, jan 1997. doi: 10.1099/00221287-143-1-179.

35. Oskar Hallatschek, Pascal Hersen, Sharad Ramanathan, and David R Nelson. Genetic drift at expanding frontiers promotes gene segregation. PNAS, 104(50):19926–30, dec 2007. ISSN 1091-6490. doi: 10.1073/pnas.0710150104.

36. Yoshifumi Imamura et al. Azithromycin exhibits bactericidal effects on Pseudomonas aeruginosa through interaction with the outer membrane. Antimicrobial agents and chemotherapy, 49(4):1377–80, apr 2005. doi: 10.1128/AAC.49.4.1377-1380.2005.

37. Frank Oechslin, Philippe Piccardi, Stefano Mancini, Jérôme Gabard, Philippe Moreillon, José M. Entenza, Gregory Resch, and Yok-Ai Que. Synergistic interaction between phage therapy and antibiotics clears *Pseudomonas aeruginosa* infection in endocarditis and reduces virulence. Journal of Infectious Diseases, 67:jiw632, dec 2016. doi: 10.1093/infdis/jiw632.

38. Shuai Le et al. Chromosomal DNA deletion confers phage resistance to Pseudomonas aeruginosa. Scientific Reports, 4(1):4738, may 2015. doi: 10.1038/srep04738.

39. S. J. Schrag and J. E. Mittler. Host-Parasite Coexistence: The Role of Spatial Refuges in Stabilizing Bacteria-Phage Interactions. The American Naturalist, 148(2):348–377, aug 1996. doi: 10.1086/285929.

40. James J Bull, Kelly A Christensen, Carly Scott, Benjamin R Jack, Cameron J Crandall, and Stephen M Krone. Phage-Bacterial Dynamics with Spatial Structure: Self Organization around Phage Sinks Can Promote Increased Cell Densities. Antibiotics (Basel, Switzerland), 7(1), jan 2018. doi: 10.3390/antibiotics7010008.

41. Lucia Vidakovic, Praveen K. Singh, Raimo Hartmann, Carey D. Nadell, and Knut Drescher. Dynamic biofilm architecture confers individual and collective mechanisms of viral protection. Nature Microbiology, 3(1):26–31, jan 2018. ISSN 2058-5276. doi: 10.1038/ s41564-017-0050-1.

42. E Chapman-McQuiston and X L Wu. Stochastic receptor expression allows sensitive bacteria to evade phage attack. Part I: experiments. Biophysical journal, 94(11):4525–36, jun 2008. doi: 10.1529/biophysj.107.120212.

43. Silja Heilmann, Kim Sneppen, and Sandeep Krishna. Coexistence of phage and bacteria on the boundary of self-organized refuges. Proceedings of the National Academy of Sciences, 109(31):12828–12833, jul 2012. ISSN 0027-8424. doi: 10.1073/pnas.1200771109.

44. Matthew Simmons, Knut Drescher, Carey D Nadell, and Vanni Bucci. Phage mobility is a core determinant of phage-bacteria coexistence in biofilms. The ISME Journal, 12(2): 531–543, feb 2018. doi: 10.1038/ismej.2017.190.

45. C P Ricciuti. Host-virus interactions in Escherichia coli: effect of stationary phase on viral release from MS2-infected bacteria. Journal of virology, 10(1):162–5, jul 1972.

46. A. M. Haywood. Lysis of RNA Phage-infected Cells depends upon Culture Conditions. Journal of General Virology, 22(3):431–435, mar 1974. doi: 10.1099/0022-1317-22-3-431.

47. M. Middelboe. Bacterial Growth Rate and Marine Virus-Host Dynamics. Microbial Ecology, 40(2):114–124. doi: 10.1007/s002480000050.

48. Sanna Sillankorva, Rosàrio Oliveira, Maria João Vieira, Ian Sutherland, and Joana Azeredo. Pseudomonas fluorescens infection by bacteriophage ΦS1: the influence of temperature, host growth phase and media. FEMS Microbiology Letters, 241(1):13–20, dec 2004. ISSN 03781097. doi: 10.1016/j.femsle.2004.06.058.

49. Robin A. Sorg et al. Collective Resistance in Microbial Communities by Intracellular Antibiotic Deactivation. PLOS Biology, 14(12):e2000631, dec 2016. doi: 10.1371/journal.pbio. 2000631.

50. Marjon G J de Vos, Marcin Zagorski, Alan McNally, and Tobias Bollenbach. Interaction networks, ecological stability, and collective antibiotic tolerance in polymicrobial infections. Proceedings of the National Academy of Sciences of the United States of America, 114(40): 10666–10671, oct 2017. doi: 10.1073/pnas.1713372114.

51. Eugene Anatoly Yurtsev, Arolyn Conwill, and Jeff Gore. Oscillatory dynamics in a bacterial cross-protection mutualism. Proceedings of the National Academy of Sciences, 113(22): 6236–6241, may 2016. doi: 10.1073/pnas.1523317113.

52. Isabel Frost et al. Cooperation, competition and antibiotic resistance in bacterial colonies. The ISME Journal, 12(6):1582–1593, jun 2018. ISSN 1751-7362. doi: 10.1038/ s41396-018-0090-4.

53. Carsten Matz and Staffan Kjelleberg. Off the hook - how bacteria survive protozoan grazing. Trends in Microbiology, 13(7):302–307, jul 2005. doi: 10.1016/J.TIM.2005.05.009.

54. Elhanan Tzipilevich, Michal Habusha, and Sigal Ben-Yehuda. Acquisition of Phage Sensitivity by Bacteria through Exchange of Phage Receptors. Cell, 168(1–2):186–199.e12, jan 2017. doi: 10.1016/J.CELL.2016.12.003.

55. Pavel Payne, Lukas Geyrhofer, Nicholas H Barton, and Jonathan P Bollback. CRISPR-based herd immunity can limit phage epidemics in bacterial populations. eLife, 7, mar 2018. doi: 10.7554/eLife.32035.

56. Holly S. Schrader, John O. Schrader, Jeremy J. Walker, Thomas A. Wolf, Kenneth W. Nickerson, and Tyler A. Kokjohn. Bacteriophage infection and multiplication occur in *Pseudomonas aeruginosa* starved for 5 years. Canadian Journal of Microbiology, 43(12):1157–1163, dec 1997. doi: 10.1139/m97-164.

57. Nawras Ghanem, Bärbel Kiesel, René Kallies, Hauke Harms, Antonis Chatzinotas, and Lukas Y. Wick. Marine Phages As Tracers: Effects of Size, Morphology, and Physico-Chemical Surface Properties on Transport in a Porous Medium. Environmental Science & Technology, 50(23):12816–12824, dec 2016. doi: 10.1021/acs.est.6b04236.

58. M. J. I. Muller, B. I. Neugeboren, D. R. Nelson, and A. W. Murray. Genetic drift opposes mutualism during spatial population expansion. Proceedings of the National Academy of Sciences, 111(3):1037–1042, 2014. ISSN 0027-8424. doi: 10.1073/pnas.1313285111.

59. Babak Momeni, Kristen A Brileya, Matthew W Fields, and Wenying Shou. Strong interpopulation cooperation leads to partner intermixing in microbial communities. eLife, 2: e00230, jan 2013. ISSN 2050-084X. doi: 10.7554/eLife.00230.

60. Britt Koskella and Michael A. Brockhurst. Bacteria-phage coevolution as a driver of ecological and evolutionary processes in microbial communities. FEMS Microbiology Reviews, 38 (5):916–931, 2014. ISSN 1574-6976. doi: 10.1111/1574-6976.12072.

61. Andrew M Kropinski, Lora Chan, Ken Jarrell, and F H Milazzo. The nature of Pseudomonas aeruginosa strain PAO bacteriophage receptors. Canadian Journal of Microbiology, 23(6): 653–658, jun 1977. ISSN 0008-4166. doi: 10.1139/m77-098.

62. Robert B Lindberg and Ruth L Latta. Phage typing of Pseudomonas aeruginosa: Clinical and epidemiologic considerations. Journal of Infectious Diseases, 130:S33–S42, 1974.

63. M. Brown and D. G. Lowe. Automatic panoramic image stitching using invariant features. International Journal of Computer Vision, 74(1):59–73, 2007.

