## Supplementary figures for "Phage efficacy in infecting dual-strain biofilms of *Pseudomonas aeruginosa*"

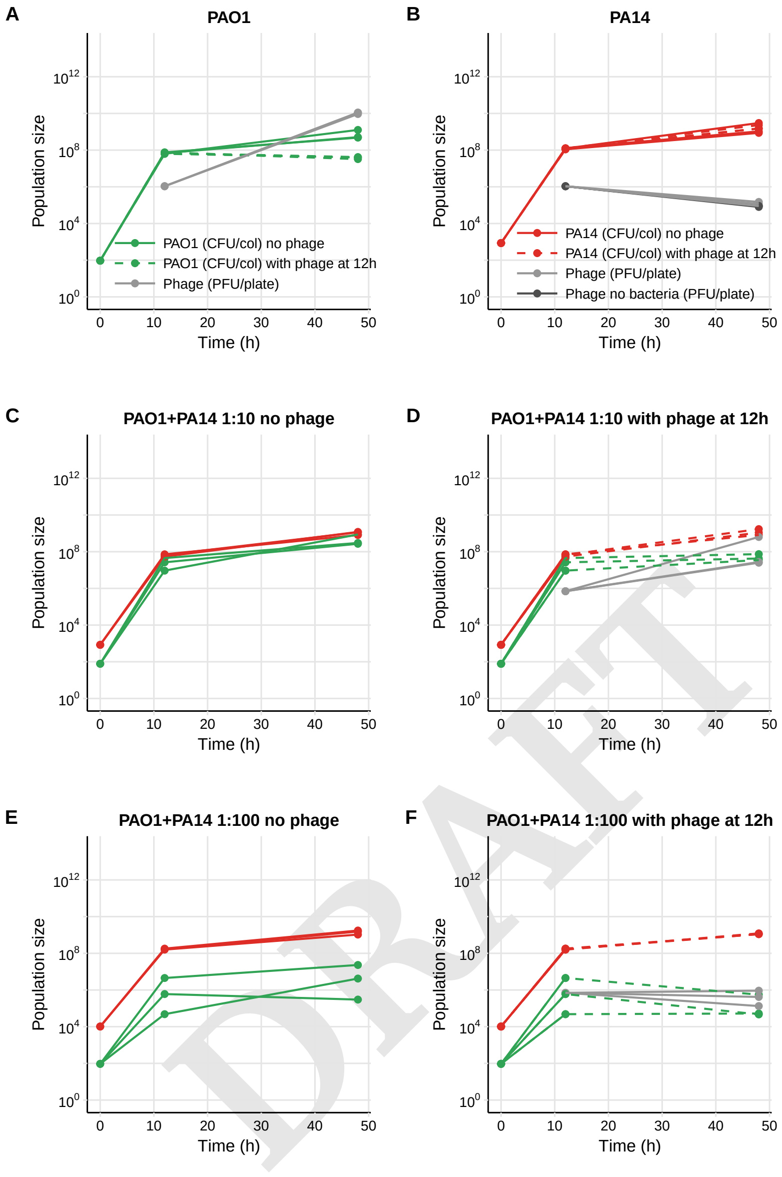


**Fig. S1.** Growth curves corresponding to data in Fig. 2C-F. (A) Growth curve of PAO1 with and without phage. PAO1 cease to grow following phage exposure. (B) PA14 with and without phage. No difference was observed following phage treatment, and phage decreased as in a control experiment with no bacteria. (C) Co-culture colony of the two strains at a 1:10 ratio, in the absence of phage. (D) Co-culture colony of the two strains at a 1:10 ratio, with phage, PAO1 again ceases to grow significantly. (E) Same as (C) but with a 1:100 ratio between the two strains (higher initial population size of PA14). Due to the increased competition, PAO1 cannot grow as well. (F) Same as (D) but with a 1:100 ratio between the two strains. Again, due to increased competition, there are fewer PAO1 cells, so phage cannot replicate as much. This leads to the lower overall phage population size as shown in Fig. 2F.

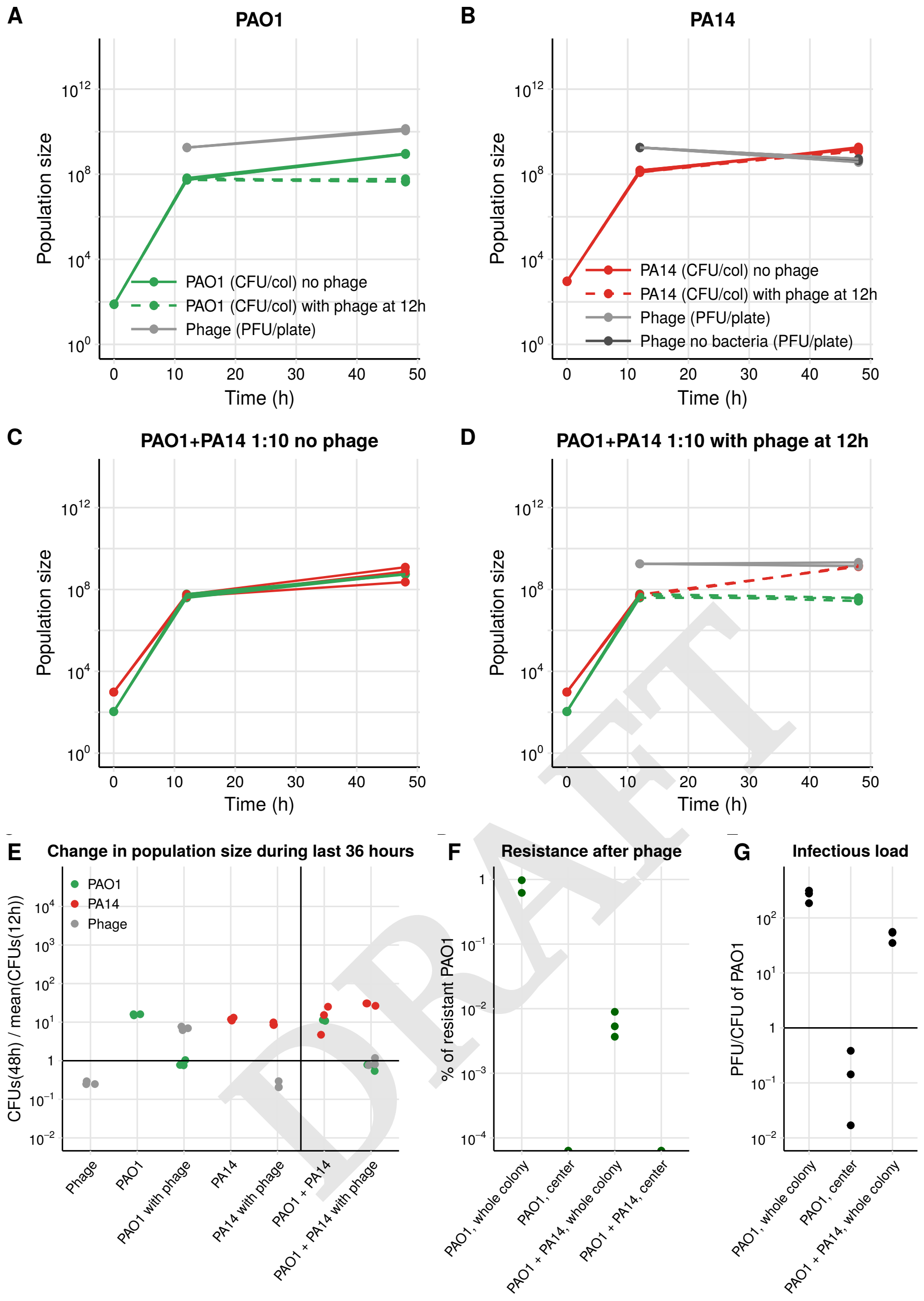


**Fig. S2.** Growth curves corresponding to data in Fig. S1 and 2C-F, but with an initial phage inoculum of ~$10^9$/ml. (A) Growth curve of PAO1 with and without phage. PAO1 cease to grow following phage exposure. (B) PA14 with and without phage. No difference was observed following phage treatment, and phage decreased as in a control experiment with no bacteria. (C) Co-culture colony of the two strains at a 1:10 ratio, in the absence of phage. (D) Co-culture colony of the two strains at a 1:10 ratio, with phage, PAO1 again ceases to grow significantly, but this time the phage do not increase significantly, probably because the population size is already very large. (E) Corresponding panel to Fig. 2C. The main difference is the lack of increase of the phage population. (F) Corresponding panel to Fig. 2D. Resistance to the phage was significantly higher in the mono-culture colonies here, but the center still only contained sensitive cells. Furthermore, although it appears that all cells were resistant, we picked 19 of the CFUs from the phage-free agar plates to test for phage resistance (see Methods), and found that at least 5 out of 19 proved to still be sensitive to the phage. In the co-culture colonies, we now detect resistance at the edges. Presumably, more phage poses a greater selection pressure leading to increased resistance. However, it is still clear that resistance is less likely in the mixed colonies. (G) To determine whether phage could diffuse into the colonies, we touched the centers with an inoculation loop and counted the PAO1 CFUs and phage PFUs. Results are comparable to Fig. 2E.

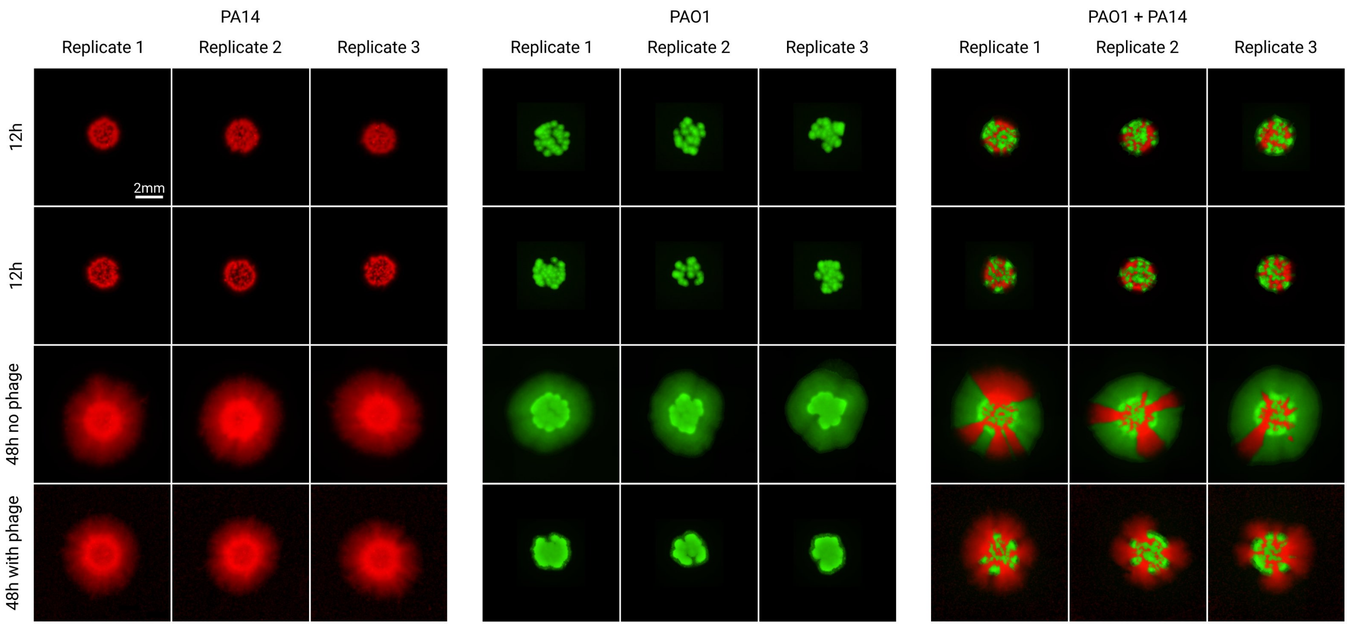


**Fig. S3.** Fluorescence microscopy images of colonies at 12 and 48 hours in a preliminary experiment similar to those shown in Fig. 2B. PA14 are tagged with mCherry (red) and PAO1 with GFP (green). The first row shows the colonies that were transferred to agar without phage, while the second row are the colonies that were later exposed to phage. The first and second row therefore correspond to the images in the third and forth row, respectively.

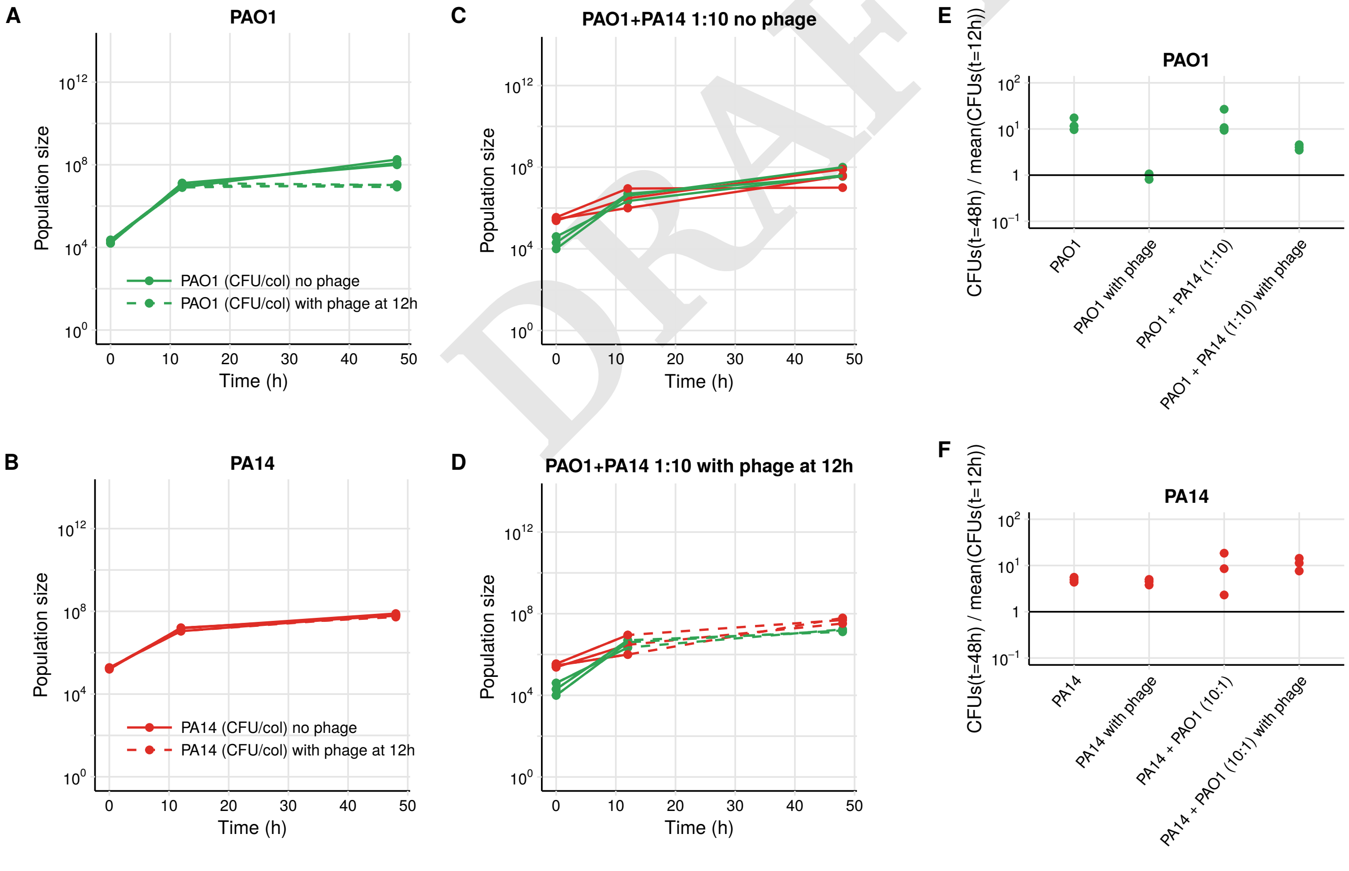


DRAFT

**Fig. S4.** Growth curves corresponding to images in Fig. S3. (A) Growth curve of PAO1 with and without phage. PAO1 cease to grow following phage exposure. (B) PA14 with and without phage. No difference was observed following phage treatment. We did not measure phage population sizes in this experiment. (C) Co-culture colony of the two strains at a 1:10 ratio, in the absence of phage. (D) Co-culture colony of the two strains at a 1:10 ratio, with phage. (E-F) Corresponding panel to Fig. 2C but for (E) PAO1 and (F) PA14.

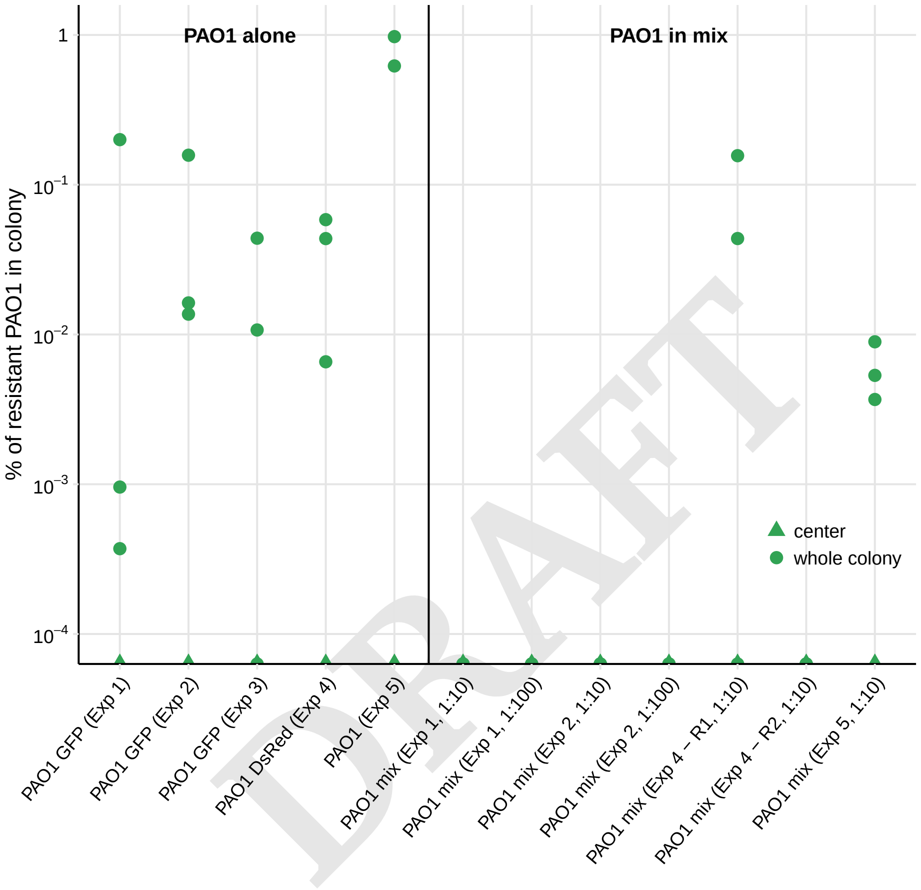

DRAFT

**Fig. S5.** Similar to Fig. 2D, but including data from all experiments for comparison. Experiment 1 corresponds to the data in Fig. 2B and S6, experiment 2 to data in Fig. 2C-F and S1, experiment 3 was a repeat of experiment 2 (data not shown), experiment 4 to data shown in Fig. S15, and experiment 5 was with higher phage dose, corresponding to Fig. S2. We never found resistance in the center of any colonies. We only found resistance in mixed colonies either if the resistant strain was completely outcompeted by the sensitive one, in which case the colony behaved as a mono-culture (Exp 4 - resistant strain R1); or if the initial dose of phage as 3 orders of magnitude higher, leading to increased selection pressure (Exp 5).

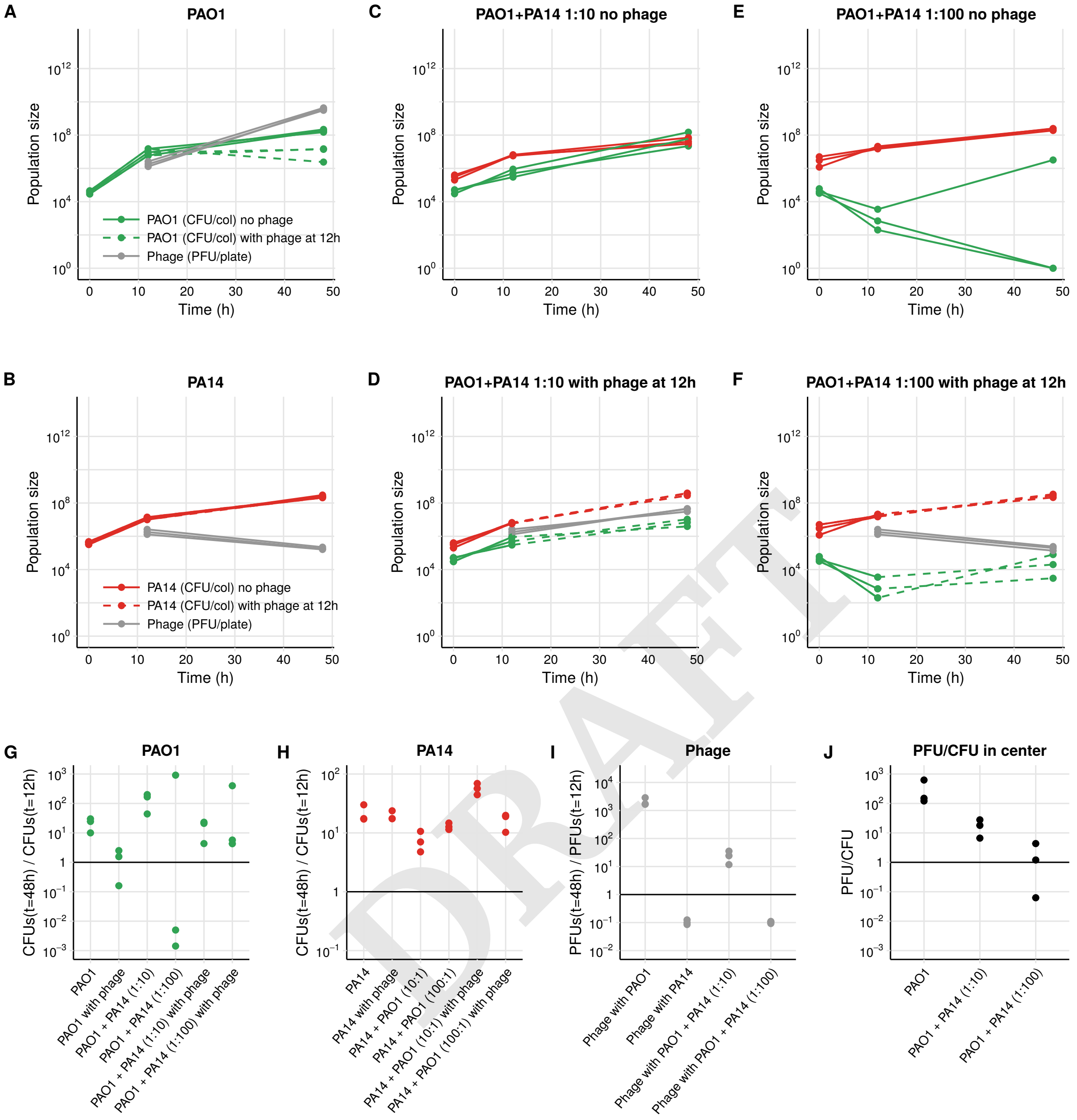


DRAFT

**Fig. S6.** Growth curves corresponding to data in Fig. 2B. Here, PA14-mCherry was used, making it harder to quantify population sizes compared to Fig. S1. (A) Growth curve of PAO1 with and without phage. PAO1 cease to grow following phage exposure. (B) PA14-mCherry with and without phage. No difference was observed following phage treatment, and phage decreased as in a control experiment with no bacteria. (C) Co-culture colony of the two strains at a 1:10 ratio, in the absence of phage. (D) Co-culture colony of the two strains at a 1:10 ratio, with phage, PAO1 again ceases to grow significantly. (E) Same as (C) but with a 1:100 ratio between the two strains (higher initial population size of PA14-mCherry). Due to the increased competition, PAO1 cannot grow as well. (F) Same as (D) but with a 1:100 ratio between the two strains. Again, due to increased competition, there are fewer PAO1 cells, so phage cannot replicate as much. This leads to the lower overall phage population size as shown in Fig. 2F. (G) The ratio of population sizes of PAO1 at 48 and 12 hours in the different colonies. (H) The ratio of population sizes of PA14-mCherry at 48 and 12 hours. (I) The ratio of population sizes of of phage at 48 and 12 hours. (J) To determine whether phage could diffuse into the colonies, we touched the centers with an inoculation loop and counted the PAO1 CFUs and phage PFUs. Results are comparable to Fig. 2E.

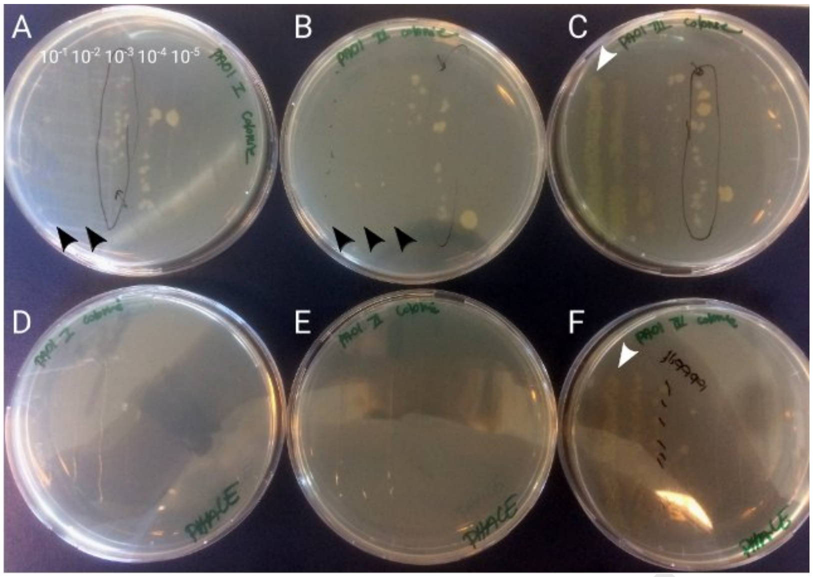


**Fig. S7.** Dilutions performed to count CFUs. The petridishes in the top row contained agar only, while those in the bottom row were saturated with phage. In each petridish, parallel lines were plated, each from a dilution of the sampled PAO1 mono-culture colony. The three columns show three replicates. In the top row, the black arrowheads show dilutions at which nothing grew, presumably because cells containing phage lysed and infected their neighbours, eliminating all bacteria. As the cultures were diluted, individual uninfected bacteria become visible as they are able to form CFUs. It is these CFUs that could potentially cause problems from a therapeutic point of view: if such uninfected cells are dispersed and land on a new surface far from other infected cells, they could start a new bacterial infection. In the third replicate on the right, there are also resistant cells, which form CFUs both at high dilutions (white arrowhead) in the top row and in the bottom row, where the plates were saturated with phage. The other two replicates contained no resistant bacteria.

DRAFT

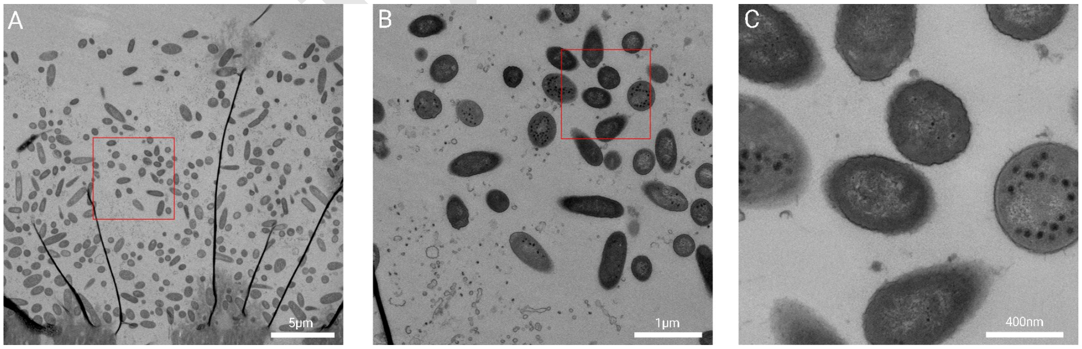


**Fig. S8.** Transmission electron microscopy image of the colony. (A) A slice through a colony, with the filter visible at the bottom, and at the top the empty zone above the colony. We cannot be certain of the location of the image within the colony. The dark black lines are folds that occurred as an artifact during sample preparation. The red box shows the location of panel (B). (B) Phage start to become visible both inside and around the cells. Note that cell debris from lysed cells is also visible in the bottom left corner or the image. The red box shows the location of panel (C). (C) example of some cells containing assembled phage, and others that are free of phage. Again, cells containing phage have not lysed, suggesting delayed lysis, but not “pseudolysogeny”, where we would not expect to see assembled phage capsids.

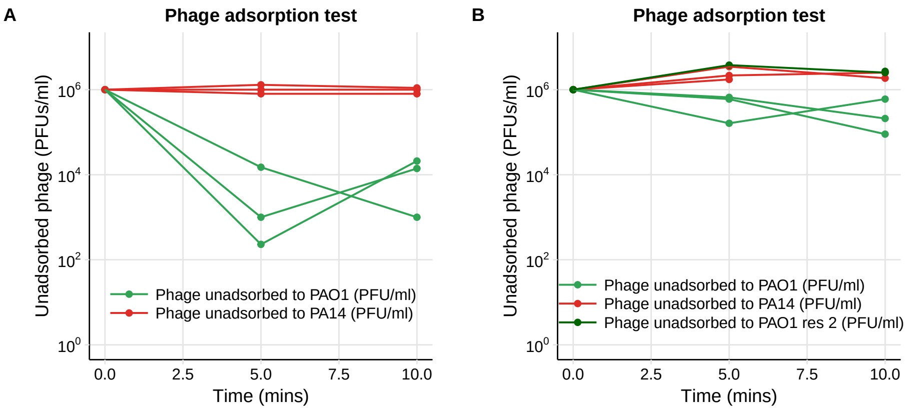


**Fig. S9.** Adsorption test. Different strains were mixed with phage for 5 or 10 minutes, then centrifuged and the supernatant used to count PFUs. If not attachment occurred, the phage population size remained the same, whereas phage attachment resulted in a decrease in recovered population size. (A) Comparing PA14 and PAO1. (B) Repeat of the previous assay, in addition to PAO1 resistant strain 2. Initial phage numbers are theoretical based on the preparation of the phage stocks.

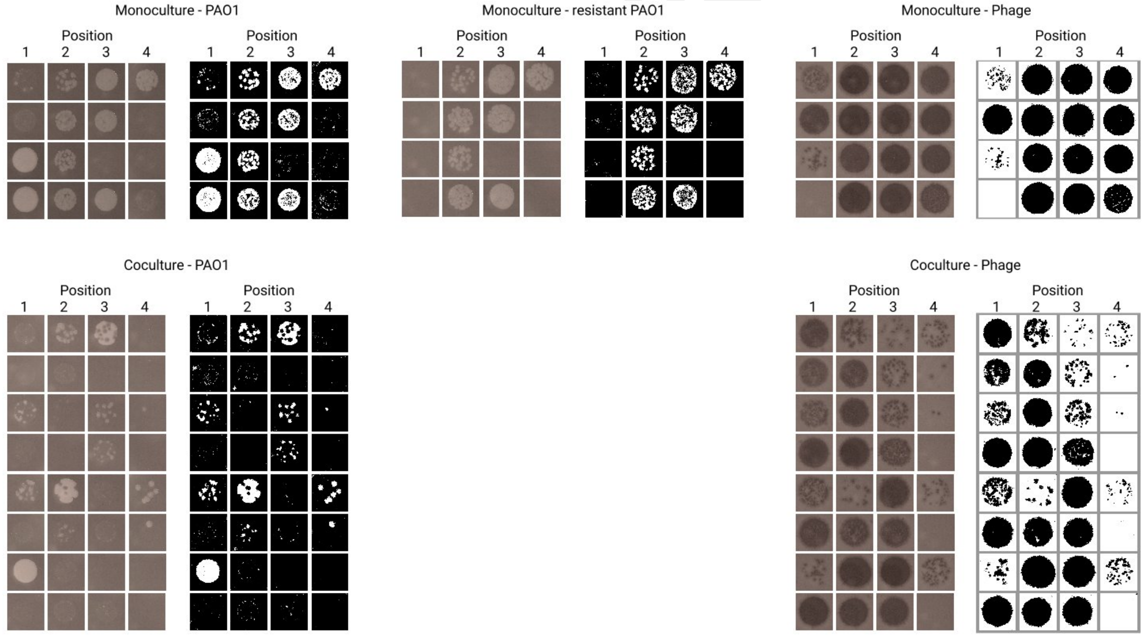


**Fig. S10.** Dataset used to generate plot in Fig. S12. Positions correspond approximately to those touched with the toothpick as shown in Fig. 3A. On the left we show all original images, and to the right the thresholded images (see Methods). These thresholded images are then used to compute the density of bacteria and phage. Images for resistant PAO1 in the co-culture condition are not shown since nothing grew.

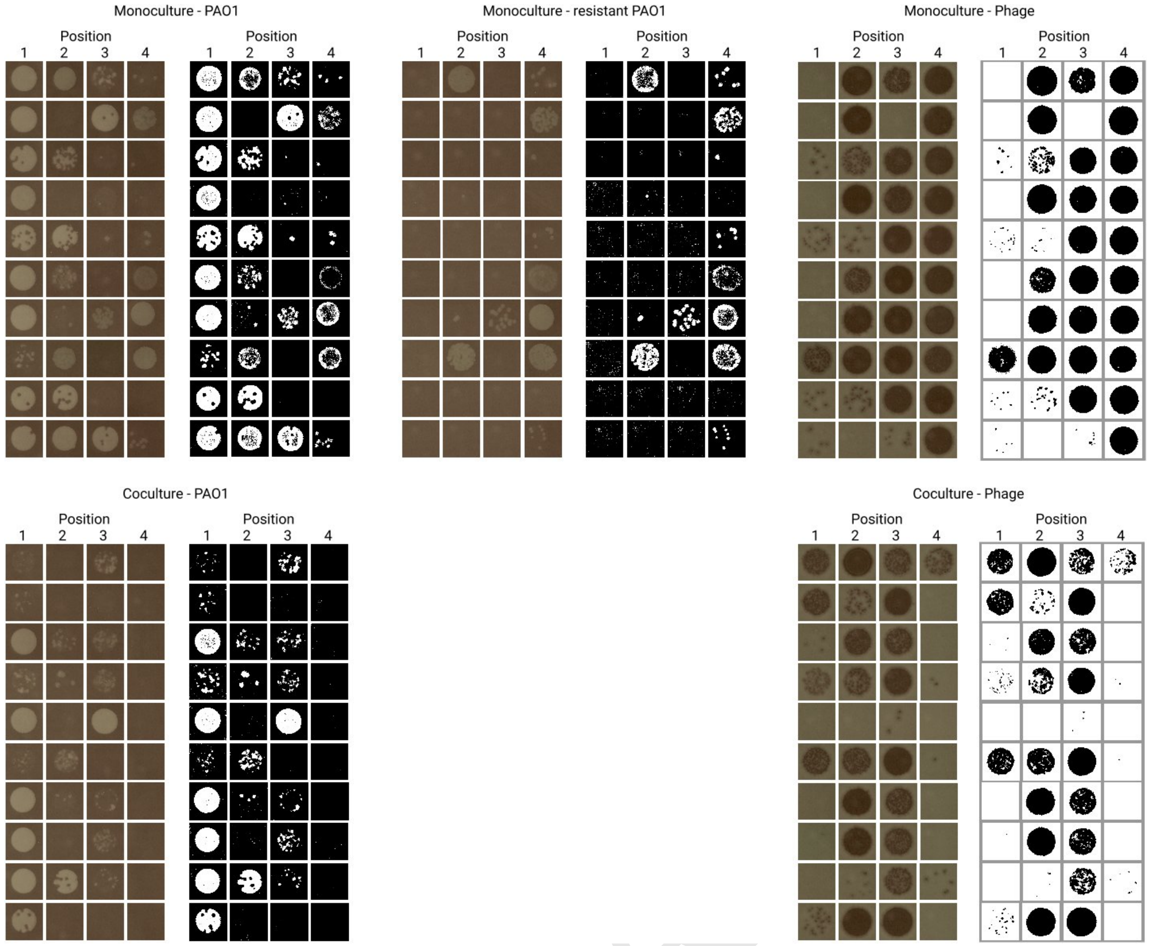


**Fig. S11.** Dataset used to generate plot in Fig. 3. Positions correspond approximately to those touched with the toothpick as shown in Fig. 3A. On the left we show all original images, and to the right the thresholded images (see Methods). These thresholded images are then used to compute the density of bacteria and phage. Images for resistant PAO1 in the co-culture condition are not shown since nothing grew.

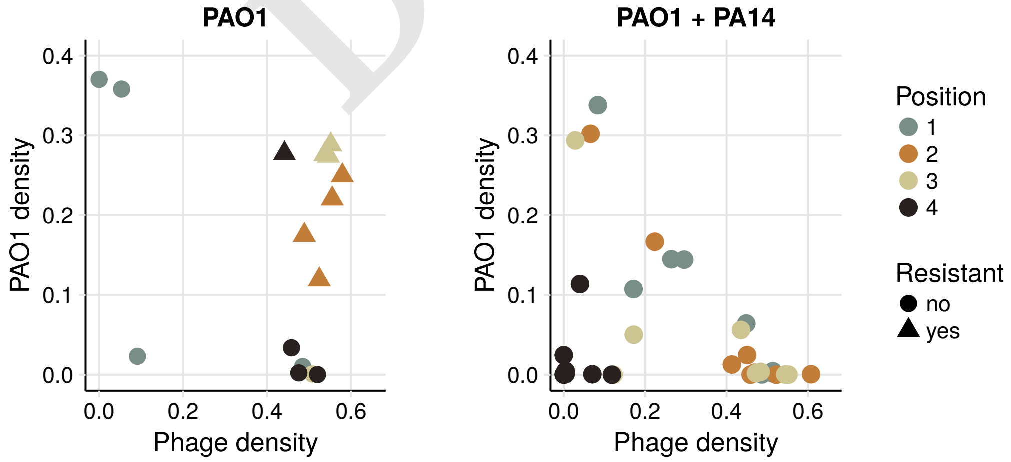


**Fig. S12.** Sampling colonies to determine co-occurrence patterns of phage and bacteria (repeat of experiment shown in Fig. 3). Each dot or triangle corresponds to a sample in one position in one colony. The left and right panels show samples taken from 4 PAO1 (16 points) and 8 mixed colonies (32 points). Resistance was determined by similarly thresholding images of drops grown on LB agar with gentamicin and saturated with ~$10^{10}$ phage (see Fig. S10 for the full data set). The different colors represent the positions sampled as shown in Fig. 3A.

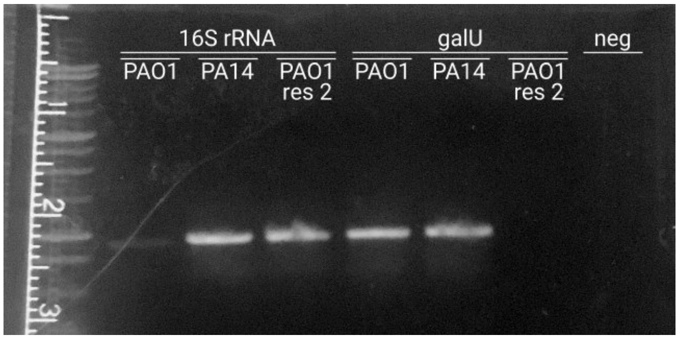


**Fig. S13.** Photo of gel electrophoresis where we ran the PCR product of the amplified 16S gene of PAO1, PA14 and PAO1 resistant strain 2, and of the amplified galU gene of the same three strains. This shows that the galU gene is absent in the resistant strain.

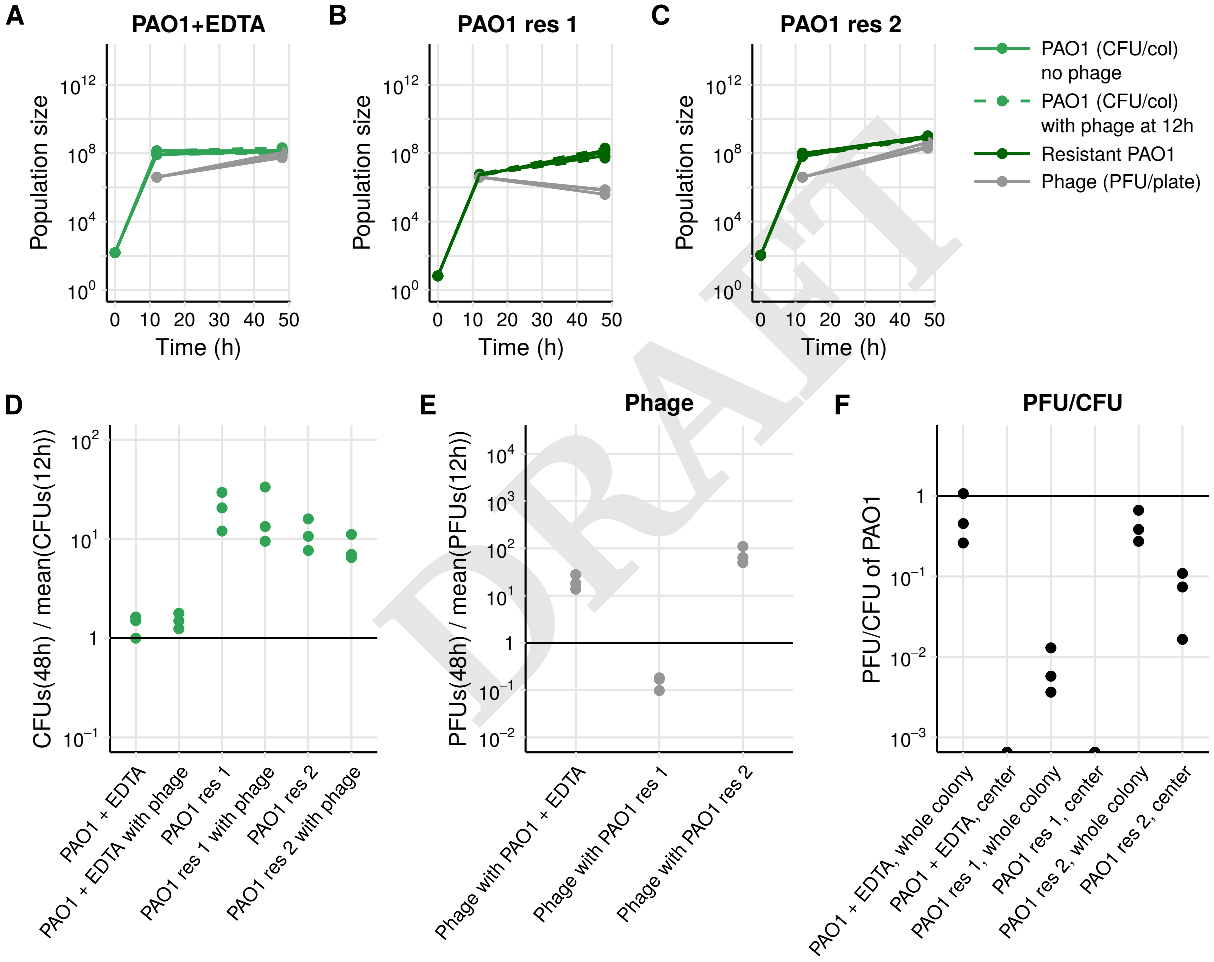


DRAFT

**Fig. S14.** Experiment with mono-cutlures of the wildtype PAO1 growing on EDTA and two resistant PAO1 isolates growing on LB. (A) Growth curve of PAO1 on EDTA with and without phage. Even though the phage replicates somewhat, there is mostly no growth or death. (B) The first resistant isolate, studied in more detail in the main text, is completely resistant, such that the phage decrease as in Fig. 2C with phage alone, or with PA14.(C) The second resistant strain is only partially resistant (approx. 1 in 10 CFUs were able to form colonies on phage-saturated agar plates), such that the phage can still replicate. (D) The ratio of population sizes of bacteria at 48 and 12 hours. (E) The ratio of population sizes of phage at 48 and 12 hours. (F) To determine whether phage could diffuse into the colonies, we touched the centers with an inoculation loop and counted the PAO1 CFUs and phage PFUs. No phage were detected in the centers of colonies growing on EDTA or completely resistant colonies. All colonies contained fewer phage than in the data shown in Fig. 2E.

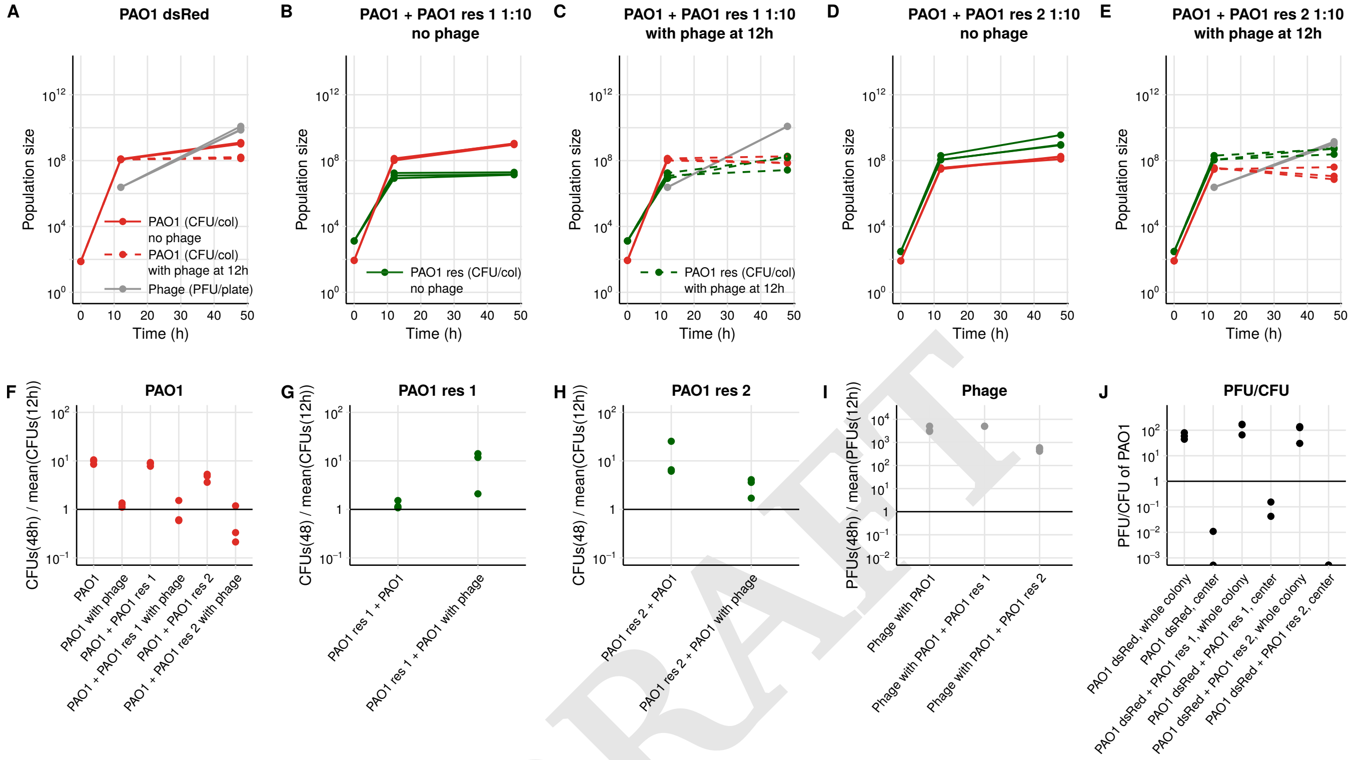


DRAFT

**Fig. S15.** Experiment 4 where we replaced PA14 with one of two resistant PAO1 isolates (resistant strain 1 (galU mutant, Fig. S13) which is fully resistant, and resistant strain 2 which is partially resistant (Fig. S14B, C)). To distinguish the two strains when in co-culture colonies, we used a wildtype PAO1 strain with a DsRed tag. The resistant strains were labelled with GFP. (A) Growth curve of PAO1-DsRed with and without phage, comparable to Fig. S1. (B) Growth curve of PAO1-DsRed when in co-culture with resistant strain 1, with no phage. The fitness cost to becoming completely resistant to phage is visible in its slow growth. (C) Same as (B) but with phage. Here, the phage infects and replicates similar to the case where wildtype PAO1 is in mono-culture. (D) Same as (B) but with resistant strain 2. (E) Same as (D) but with phage. (F) The ratio of population sizes of PAO1-DssRed at 48 and 12 hours in the different colonies. (G) The ratio of population sizes of PAO1 resistant strain 1 at 48 and 12 hours. (H) The ratio of population sizes of PAO1 resistant strain 2 at 48 and 12 hours. (I) The ratio of population sizes of phage at 48 and 12 hours. (J) To determine whether phage could diffuse into the colonies, we touched the centers with an inoculation loop and counted the PAO1 CFUs and phage PFUs. All three treatments behave similar to PAO1-GFP mono-cultures (Fig. 2E), PAO1-DsRed because it is isogenic, with resistant strain 1 because it grows so little, leaving PAO1-DsRed to dominate as in mono-culture, and resistant strain 2 because it can also be infected, so there are many phage.
